# Mechanical canalization of chiral gut morphogenesis

**DOI:** 10.64898/2026.11.17.766015

**Authors:** Nicolas Romeo, Chris Anto, Avi Strok, Emily L. Hendricks, Natalia A. Tamarina, Andrew Fee, Fridtjof Brauns, Noah P. Mitchell

## Abstract

Chiral organs such as the gut develop complex, stereotyped shapes, despite variability in the molecular and cellular processes that build them. How cell activity specifies tissue-scale handedness and reproducibly shapes whole organs remains unclear. Here, we use the embryonic *Drosophila* midgut to understand how a symmetric tubular organ reliably coils into a complex loop of alternating handedness, without any known left–right asymmetric gene expression. Through multi-view light-sheet imaging, 3D geometric analysis, and physical modeling, we show that midgut handedness is specified by epithelial cells in a neighboring organ, relayed across all three germ layers, and finally amplified and canalized by the mechanics of confined gut elongation. Myosin 1C expression in the hindgut epithelium controls midgut handedness up to hundreds of microns away, transmitting chiral bias to the midgut through migratory cells that form the longitudinal muscle. As the midgut elongates, a buckling instability then amplifies early left–right asymmetric constrictions, and confinement funnels the organ into a stereotyped coil. Tissue mechanics thus canalizes organ shape into one of two chiral forms, selected by an inter-organ cellular relay.

---

Many internal organs adopt complex, stereotyped three-dimensional (3D) shapes of a specific handedness: the vertebrate heart tube loops to the right [1], and the gut coils into a reproducible arrangement of loops [2, 3]. These chiral configurations allow elongated organs such as the gut to pack within the body and function correctly [4–7]. Their shape must therefore be both reproducible and of the specified handedness relative to the rest of the body [8]. Such reproducibility despite variability in underlying processes, termed canalization [9–13], has mainly been understood through regulatory mechanisms that stabilize cell states and developmental trajectories [14–18], yet organ shape is ultimately generated by mechanical forces [19], which can themselves suppress local variability during morphogenesis [20–26]. How mechanics and molecular cues together make complex 3D organ shapes reproducible remains largely open.

Left–right asymmetry can arise in two ways. In vertebrates and several invertebrates, asymmetric expression of genes such as *Nodal* instructs organ laterality [27– 31]. Independently, the chiral organization of the cytoskeleton generates intrinsic handedness within individual cells. This cell-intrinsic mechanism is increasingly recognized as a source of organ asymmetry in vertebrates and invertebrates alike [32–36]. The *Drosophila melanogaster* digestive tract makes it possible to isolate how tissues translate cell-scale chirality into chiral organ shapes: these organs break left–right symmetry without any known left–right asymmetric gene expression. Its embryonic midgut, for instance, folds from a symmetric tube into four chambers arranged in a stereotyped 3D coil. The handedness of this coil is linked to unconventional myosins Myo1C and Myo1D [37, 38], which break symmetry at the molecular scale through chiral interactions with actin filaments [39, 40]. Yet how molecular chirality determines organ-scale handedness and how tissue mechanics shape the resulting asymmetry into a stereotyped coil remain unclear [37, 41].

Here, combining light-sheet imaging of genetically cleared embryos, 3D morphometrics, physical modeling, and genetics, we trace the origin and reproducibility of chiral midgut morphogenesis. First, migratory cells that form the midgut’s longitudinal muscle convey chirality from one organ to another: a cytoskeletal bias generated in the ectodermal hindgut is relayed by muscle precursors into the endodermal midgut, controlling initial asymmetry through left–right asymmetric constrictions. Subsequently, as the organ then elongates within the confined body cavity, a buckling instability amplifies this small bias into a stereotyped coil, canalizing the midgut into one of two stable, near mirror-image forms.

## CHIRAL COILING IS STEREOTYPED YET INVERTIBLE

The embryonic midgut develops into a stereotyped chiral coil composed of functionally distinct regions. By embryonic stage 15, two multilayered sheets have fused to form the midgut tube. The endoderm, circular visceral muscle (CVM), and longitudinal visceral muscle (LVM) arrange as three concentric layers in the closed tube (Fig. 1a-b). Three localized constrictions partition the tube into four chambers, which subsequently arrange into a chiral, alternating left–right–left–right sequence during stage 16. Remarkably, overexpression of Myosin 1C (Myo1C) in the *brachyenteron* (*byn*) domain inverts organ handedness (Fig. 1c; [37, 38, 42]).

**FIG 1.**
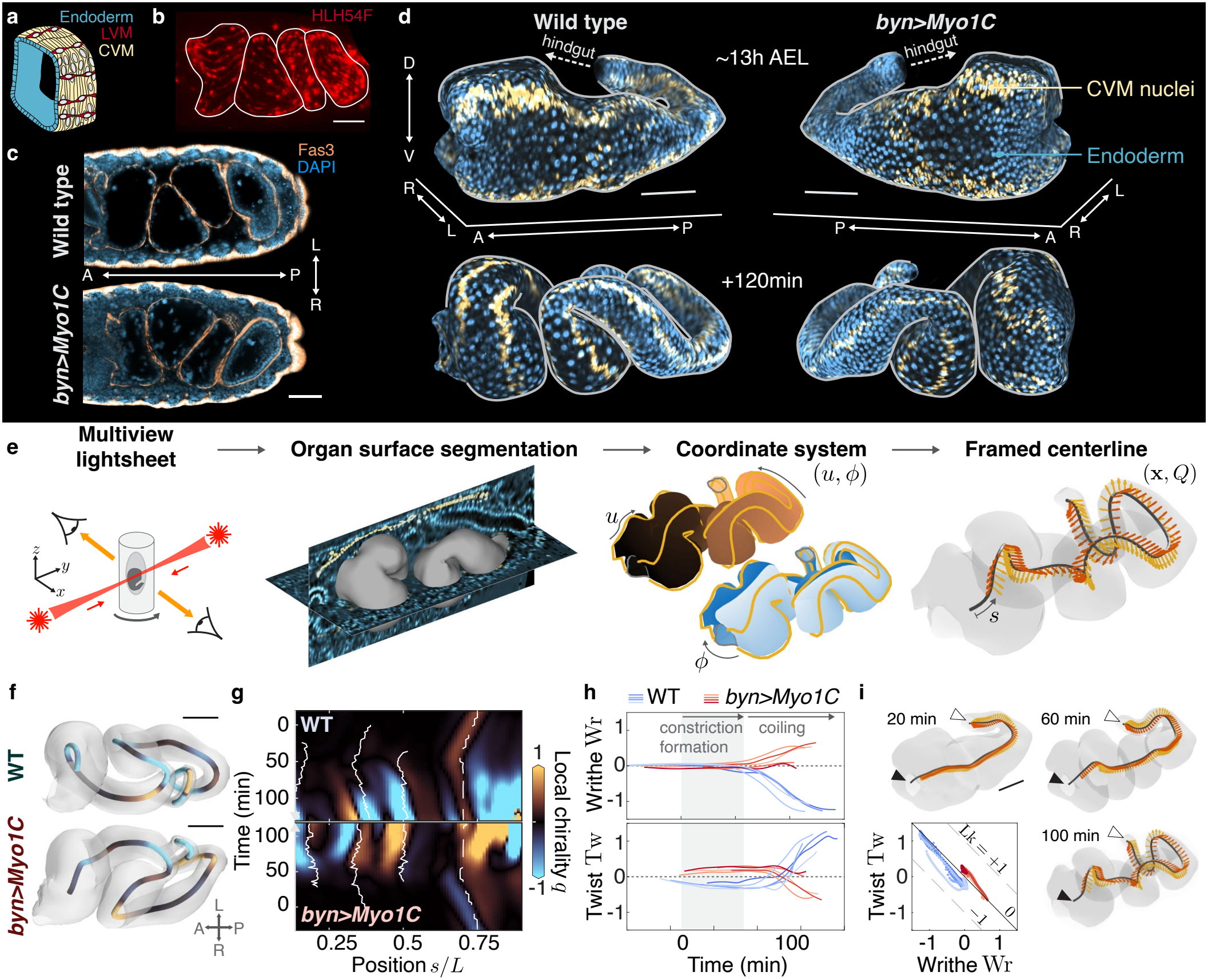
Chiral coiling of the embryonic midgut is stereotyped yet invertible. **a**. The midgut comprises an epithelium (blue) surrounded by circular (CVM, yellow; nuclei highlighted) and longitudinal (LVM, red) visceral muscle. **b**. HLH54F.LVM-RFP signal marks the LVM. **c**. Stage 16 embryos stained for visceral mesoderm (Fas3) and nuclei (DAPI), ventral views: the wild-type midgut forms a left-handed coil, mirrored by overexpressing Myo1C in the *byn* domain. **d**. About 13 h after egg laying (AEL), the midgut deforms from a tube into a coil within 2 h. **e**. From 3D movies, we extract parametrized surface meshes with annotated CVM nuclei, which define an anatomical frame for the organ centerline. **f**,**g**. Local chirality *q* along the centerline reveals that the midgut is a perverted coil with segments of alternating handedness. At the final time point, *byn>Myo1C* midguts are close to point-wise mirror images of wild type. (see Fig. S10 for all samples) Solid lines, constrictions; dashed line, posterior-most point. Time *t* is measured from the first constriction of a reference wild-type embryo, with samples aligned as in SI Sec. III. **h**. Writhe Wr and Twist Tw of wild-type and *byn>Myo1C* midguts (*n* = 5 each). Shaded band denotes the approximate period of constriction formation. **i**. Since the ends of the midgut move and rotate little, Wr + Tw remains approximately constant. Scale bars, 50 *μ*m.

To understand how this perturbation mirrors the midgut, we must identify where its chirality is generated. A natural candidate is the hindgut, which lies immediately posterior to the midgut and breaks left–right symmetry earlier. The two organs have distinct developmental origins, the hindgut being ectodermal and the midgut endodermal, and join into a continuous tube at the pylorus [43, 44]. They are nonetheless connected, both mechanically and through LVM precursors: mesenchymal myoblasts originate posteriorly and migrate anteriorly to ensheathe the midgut [45–47]. Earlier work suggested that the midgut’s handedness is determined by the hindgut’s handedness [37, 48], but the two organs invert at different frequencies in some genetic backgrounds [49, 50], challenging this hypothesis. The midgut may therefore set its handedness autonomously, or inherit a bias from the hindgut, either mechanically–for instance if the hindgut twisted the posterior end of the midgut–or through instructive signals. We first tested for a mechanical bias by following the 3D shape dynamics of the midgut, and then probed instructive cues genetically.

To follow midgut dynamics deep inside the embryo, we combine genetically induced tissue clearing [51] with multi-view confocal light-sheet microscopy [52]. Fusing views taken from several angles yields panoramic, multi-color coverage of the whole organ at isotropic resolution throughout coiling, including deep regions that single-view imaging cannot resolve (Fig. 1d,e; *n* = 5 embryos each for wild type and *byn>Myo1C*; SI Sec. III, Fig. S3). From the segmented organ surfaces [53], we extract the midgut centerline **x**(*s*), with *s* the arc length, and an anatomical orientation frame *Q*(*s*) that tracks the left–right and dorso-ventral directions using circular muscle nuclei, whose motion is tightly coupled to the endoderm [51, 54] (Fig. 1e; SI Sec. IV, Fig. S7; Supp. Video 1).

To quantify chirality along the organ, we define a local chirality *q*(*s*), the solid angle swept by the tangent over a short arc around *s*; for short arcs, it is proportional to the Frenet-Serret torsion *τ* (*s*) [55], while summed over the whole organ it gives the Writhe Wr, which measures the overall 3D coiling of the centerline and whose sign gives its handedness [56, 57] (SI Sec. V, Fig. S9, Supp. Video 2). In contrast to previous descriptions [49], and similarly to the chick gut [2, 3], the midgut is *perverted* rather than helical [58]: it consists of segments of alternating handedness (Fig. 1f,g). The local chirality of *byn>Myo1C* midguts is close to a point-wise mirror image of wild type, except in the anterior-most region, whose handedness is controlled by a distinct mechanism [41, 59–61]. After constrictions form, Wr becomes increasingly negative in wild-type midguts, and positive in *byn>Myo1C* midguts (Fig. 1h).

The Twist Tw, the number of turns of the anatomical frame around the centerline [62, 63], shows the opposite trend, so that the total Wr + Tw is approximately conserved (Fig. 1h,i). This topological constraint reflects the fact that the ends of the midgut move and rotate little: like a ribbon coiled between fixed ends, the midgut can only coil if it twists by an equal and opposite amount (formally, the Călugăreanu–White–Fuller theorem [56, 64]; SI Sec. V.3). This has two consequences. First, no net twist enters through the ends, so that, in the absence of force-generating elements outside the tissue, coiling must be driven from within the organ rather than by its boundaries [57], arguing against a mechanical role for the hindgut. Second, the constraint favors perversions: since net coiling must be compensated by twist, which is costly for any tissue that resists torsion [65], alternating handedness lets the organ coil extensively while accumulating little net Writhe [58]. Midgut coiling is thus reproducible and intrinsic to the organ.

## SYMMETRY BREAKING PRECEDES COILING

We next ask how the left–right asymmetry arises during stage 15, and how coiling proceeds during stage 16. Constrictions of the gut surface are driven by calcium-mediated contractility of the circular visceral muscle [51]. In our timeline, the three constrictions form between *t* = 0 and *t* = 50 min (SI Sec. III.3), after LVM myoblasts have migrated over the midgut during stages 12–13 [45, 46]. During stage 15, calcium activity is localized to the future constriction sites; during stage 16, it spreads over the whole organ, narrowing the entire tube (Fig. 2a,b; Fig. S4). Because the tissue surface behaves as nearly incompressible [51] (SI Sec. III.4, Fig. S6), this narrowing elongates the centerline about four-fold in both wild-type and *byn>Myo1C* embryos (Fig. 2c). The midgut elongates only slightly during constriction formation, and Writhe appears only afterwards, once the organ elongates substantially (Fig. 2c).

**FIG 2.**
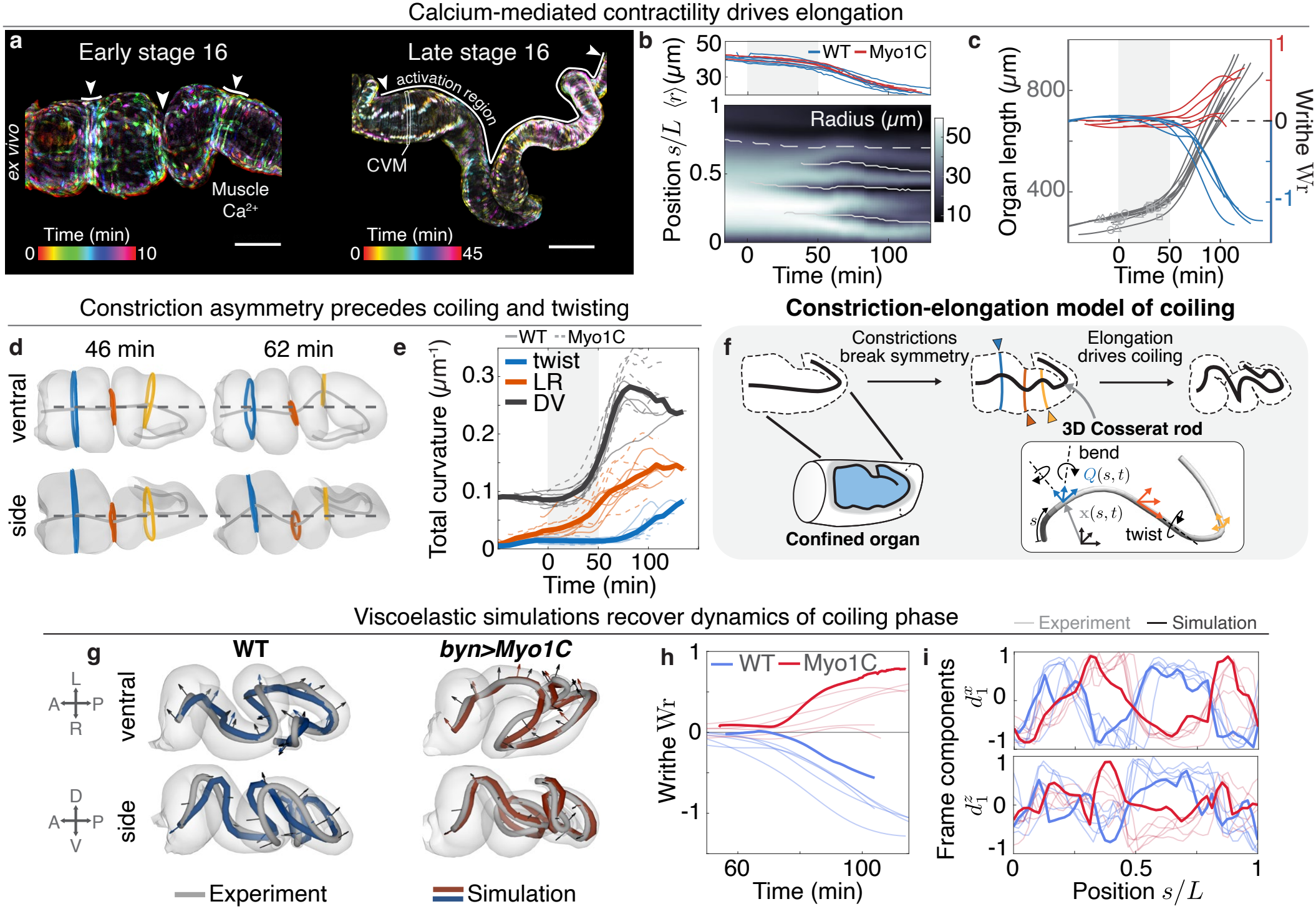
Chiral coiling results from elongation-driven buckling of an asymmetrically patterned organ. **a**. Calcium-mediated muscle activity is initially focused at the constrictions before spreading over the whole organ during stage 16, as visible in *ex vivo* midguts (*Mef2 >GCaMP8s*). **b**. Muscle activity first reduces the gut radius locally in the constriction phase, then globally in the elongation phase, as seen in the average radius (top) after constrictions form at *t* = 50 min. Solid lines, constrictions; dashed lines, posterior-most point of the midgut. Kymograph for a representative WT sample. **c**. Coiling, measured by the Writhe (blue and red, right axis), coincides with the fast phase of midgut elongation after constrictions are formed as the radius decreases (gray, left axis). Symbols indicate individual constriction formation. **d**. Constrictions initially form symmetrically with respect to the left–right and dorso-ventral axes, then break both symmetries and set the chirality pattern of the organ. **e**. During the constriction phase, left-right bending precedes twisting as measured by the integrated squared Darboux curvature, suggesting the absence of active twisting torques. **f**. The observations of **d-e** motivate a two-stage model of midgut coiling: after asymmetric constriction, elongation of the now-chiral organ within its anatomical confinement produces the coiled geometry. **g**. 3D simulations of Cosserat rods starting from the measured post-constriction shapes reproduce the observed shape dynamics in both wild-type and *byn>Myo1C* embryos. **h**. Simulations recover the Writhe dynamics. **i**. Simulations also recover the dynamics of the orientation frame, here shown for the final timepoint. See Methods for simulation parameters and definitions of measures.

Crucially, although the constrictions initially form symmetrically with respect to both the left–right (LR) and dorso-ventral (DV) axes, they are no longer aligned by the time elongation starts: the centerline breaks both symmetries and acquires a chirality pattern that matches the final coil (Fig. 2d). *byn>Myo1C* midguts display a similar DV asymmetry but the opposite LR asymmetry (Fig. S5), suggesting that the LR asymmetry of the constrictions biases the handedness of coiling.

Does this asymmetry arise from asymmetric contractility along the LR axis, or do active internal torques that wring successive segments of the organ in opposite directions [57]? By decomposing the curvature of the orientation frame into bending and twisting contributions (SI Sec. IV.4, Fig. S8), we find that the constrictions cause substantial left–right bending, whereas twist is nearly absent during constriction formation and appears only as the midgut writhes (Fig. 2e). Bending thus precedes twisting, suggesting that elongation amplifies the bends introduced by asymmetric constrictions, without requiring active torques.

To test this idea and explain how coiling proceeds once surface deformations have generated the initial asymmetry, we develop a mechanical model of the midgut. As the midgut narrows and elongates, it becomes increasingly slender and can be described by a one-dimensional (1D) framed curve. Motivated by the observation that dissected *ex vivo* embryonic midguts largely retain their shape (Fig. 2a), we model the coiling midgut as a growing viscoelastic rod (Fig. 2f; [65–68]).

The midgut is represented by its centerline and frame (**x**(*s, t*), *Q*(*s, t*)). Muscle activity sets the local elongation rate *g*(*s, t*) of the reference configuration, which we set to the measured elongation rate of the centerline. The rod resists bending and twisting, penalizing deviations of its curvature ***κ*** (related to the variation of the frame *∂*_*s*_*Q*) from a reference curvature ***κ***_0_. The reference curvature relaxes at a rate *η* toward the current curvature according to a Maxwell-like rule. We neglect spatial variations in the mechanical properties of the midgut, and assume first-order (overdamped) dynamics in the absence of evidence for more complex frictional interactions [69–71]. Further details are provided in Methods, and SI Sec. VI, Fig. S11.

We find that this setup matches experiments even with no active bending and no twisting torques, where the rod is driven only through the elongation of the centerline. The handedness of the final shape is set by the initial symmetry breaking, which is amplified by elongation-driven buckling of the rod. Simulations and experiments agree well for both centerline position and frame orientation (Fig. 2g–i; see Fig. S12, Supp. Video 3 for comparison of results across samples and snapshots of the dynamics), showing that active bending and active torques are not required for the midgut’s shape change. Notably, the overall organ shape is recovered over a wide range of dynamical parameter values (Fig. S13), suggesting that geometric effects are dominant, and reflecting the fact that a viscoelastic rod growing sufficiently fast generically preserves its initial pattern of asymmetry [72]. By recapitulating the observed coiling dynamics, the model shows that elongation-driven buckling amplifies an initial asymmetry into the mature coil, raising the question of which cells and molecular components set the direction of the initial asymmetry.

## MYOBLAST-MEDIATED RELAY OF ASYMMETRY

Modulating Myosin 1 expression in the *byn* domain is sufficient to invert midgut handedness and can rescue normal chirality of *situs inversus* mutants [37]. The *byn* domain, however, spans several tissues: *byn-GAL4* drives transient expression in the LVM precursors, strong expression in the epithelium of the pylorus and hindgut, and weak expression in the posterior-most endodermal cells of the midgut (Fig. S1).

To determine in which of these tissues Myo1C acts, we performed a GAL4/UAS overexpression screen, using the GAL4/UAS system to drive Myo1C or Myo1C-RFP expression across domains spanning the tissue layers of the midgut and hindgut (Fig. 3a). We score the handedness of each midgut at stage 16 as normal or inverted relative to wild type, and report the frequency of inversion as the fraction of inverted midguts, and report disrupted midguts as abnormal. An inversion frequency near 0 or 1 indicates a reliably specified handedness, whereas we call handedness randomized when the frequency is close to 0.5, as expected if the biasing instruction is lost. Overexpression in midgut layers–either individually or all together–did not increase the frequency of inversions, except in cases where the GAL4 driver also drove Myo1C overexpression in the hindgut epithelium (Fig. 3a). In contrast, GAL4 drivers with expression restricted to the hindgut epithelium were sufficient to invert midgut handedness. Cytoskeletal composition in the hindgut epithelium therefore instructs the direction of midgut coiling.

**FIG 3.**
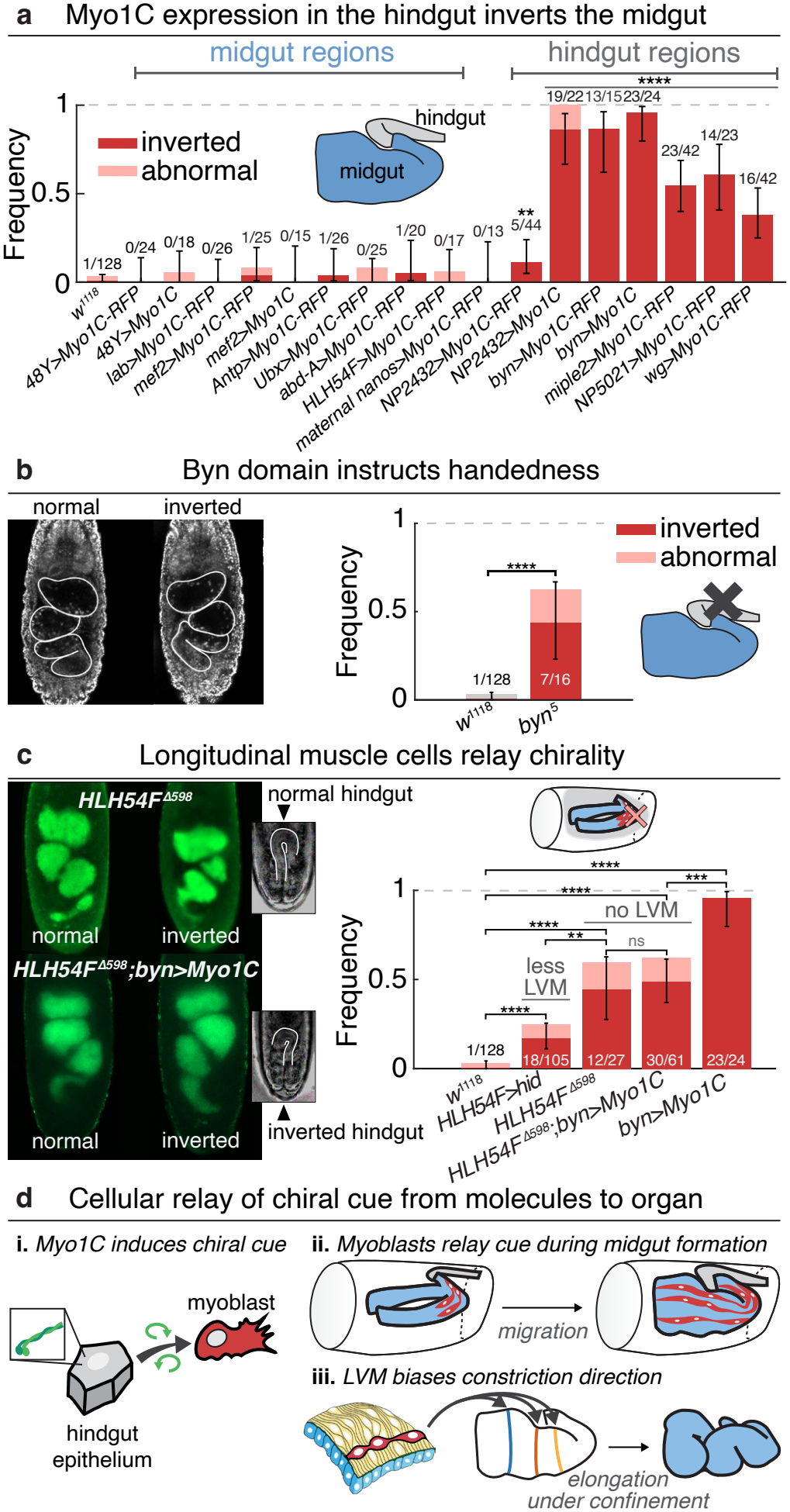
Migrating longitudinal muscle precursors relay asymmetry between two epithelia of distinct developmental origin. **a**. Myo1C overexpression in the hindgut increases midgut inversion; Myo1C overexpression restricted to midgut tissues does not. **b**. Loss of *brachyenteron* randomizes midgut handedness. **c**. Genetic ablation of the longitudinal visceral muscle (LVM) increases midgut inversion, and loss of *HLH54F*, required for LVM specification and migration, randomizes midgut chirality, regardless of whether or not Myo1C is overexpressed (*byn>Myo1C*). Hindgut chirality is normal in *HLH54F* ^Δ598^ and inverted in *HLH54F* ^Δ598^*;byn>Myo1C* embryos, showing transmission is *HLH54F* dependent. **d**. Model: chiral cytoskeletal activity in the hindgut epithelium biases neighboring myoblasts, which, once differentiated into LVM on the midgut, bias the direction of constrictions and hence of coiling. Error bars are Wilson 95% confidence in-tervals. *p*-values from a two-sided Fisher’s exact test against *w* ^1118^ (**a-c**), or as indicated (**c**). Holm-Bonferroni correction applied to hindgut and midgut drivers separately (**a**), and to all comparisons (**c**). ns, *p >* 0.05; * p*≤*0.05; ** *p ≤* 0.01; *** *p ≤* 10^*−*3^; **** *p ≤* 10^*−*4^.

Conversely, to test whether *byn*-expressing tissues are required to specify midgut handedness, we examined homozygous *byn*^5^ mutants, a reported null allele [73]. In *byn*^5^ mutants, the primordia of the hindgut epithelium undergo apoptosis [73], and LVM myoblasts largely fail to migrate and survive [46], although we find that the amount of residual LVM varies among embryos (SI Sec. II, Fig. S2). While only a small proportion of *byn*^5^ mutants advance to stage 16 (Fig. S2), those that form three constrictions were still able to coil, and these embryos’ midguts display randomized handedness (Fig. 3b). Thus, *byn*-expressing tissues are required to specify the direction of coiling, but not for coiling itself.

How, then, do cells in the posterior transmit information to midgut regions several hundred microns away along the surface? Because the LVM differentiates from mesodermally-derived myoblasts that originate posteriorly within the *byn* domain, we asked whether the LVM specifically is required to specify handedness. Genetic ablation of *HLH54F*-expressing cells, which give rise to the LVM [45], increased the frequency of inversions (*HLH54F >*hid, Fig. 3c). Similarly, embryos lacking the transcription factor HLH54F displayed randomized handedness (*HLH54F* ^Δ598^, Fig. 3c). In both conditions, the handedness and morphology of the hindgut are unaffected, so that loss of the LVM uncouples midgut handedness from that of the hindgut, in both WT and *byn>Myo1C* backgrounds (Fig. 3c, inset). Together with the conservation of Tw + Wr (Fig. 1h,i), this excludes a mechanical role for the hindgut in setting midgut handedness.

We then ask how the LVM and the hindgut epithelium interact. The LVM could relay a cytoskeletal bias originating in the hindgut epithelium, or the two tissues could act as independent controllers of midgut coiling. To distinguish these possibilities, we overexpressed Myo1C in the hindgut of embryos lacking the LVM (*HLH54F* ^Δ598^;*byn>Myo1C*, Fig. 3c). Their hindguts were inverted, but their midguts were randomized, indicating that the hindgut bias requires the LVM to reach the midgut. Notably, although the LVM is required for handedness specification, overexpressing Myo1C in the longitudinal muscle or throughout the midgut does not invert midgut handedness (*HLH54F >Myo1C-RFP, Mef2 >Myo1C* or the pan-midgut driver 48Y*>Myo1C* in Fig. 3a).

Together, these results support a model in which Myo1C activity in the hindgut epithelium generates a chiral cue that migrating myoblasts relay to the midgut. We propose that, once differentiated into LVM, these cells bias the left–right direction of the midgut constrictions, which in turn set the coiling direction (Fig. 3d), with early asymmetries amplified by buckling.

## A MECHANICAL ROUTE TO CANALIZATION

Chiral morphogenesis, here and in other chiral organs, often yields two discrete phenotypes, left- and right-handed, with few intermediates (Figs. 1f,g and 3; [36, 74– 76]). Does the buckling mechanics of our model canalizes variation in initial geometry of the midgut, beyond its robustness to material parameters?

We first simulate generic chiral rods, with initial shapes interpolating between left- and right-handed helices, growing within a tube with mechanical parameters similar to those of the midgut. Under confinement, these rods converge onto two shapes of opposite handedness (Fig. 4a,b), whereas unconfined growing rods partially preserve their initial shape [72] but do not converge (Fig. 4b, inset). As detailed in SI Sec. VIII, confinement focuses outcomes into discrete states: growth-induced buckling first favors planar shapes, which avoid twisting, but once the buckled rod fills the cavity, chiral shapes accommodate further growth with less bending than planar ones (Fig. S16; [77–79]). As growth continues, perversions emerge to limit twisting (Fig. 4c,d; SI Sec. VIII, Fig. S16, Supp. Video 4). Viscoelasticity lets the rod retain the handedness of its initial condition [72], so that a pitchfork-like bifurcation selects between the two chiral shapes—much as a slightly bowed ruler, pushed at both ends, always buckles toward its initial bow.

**FIG 4.**
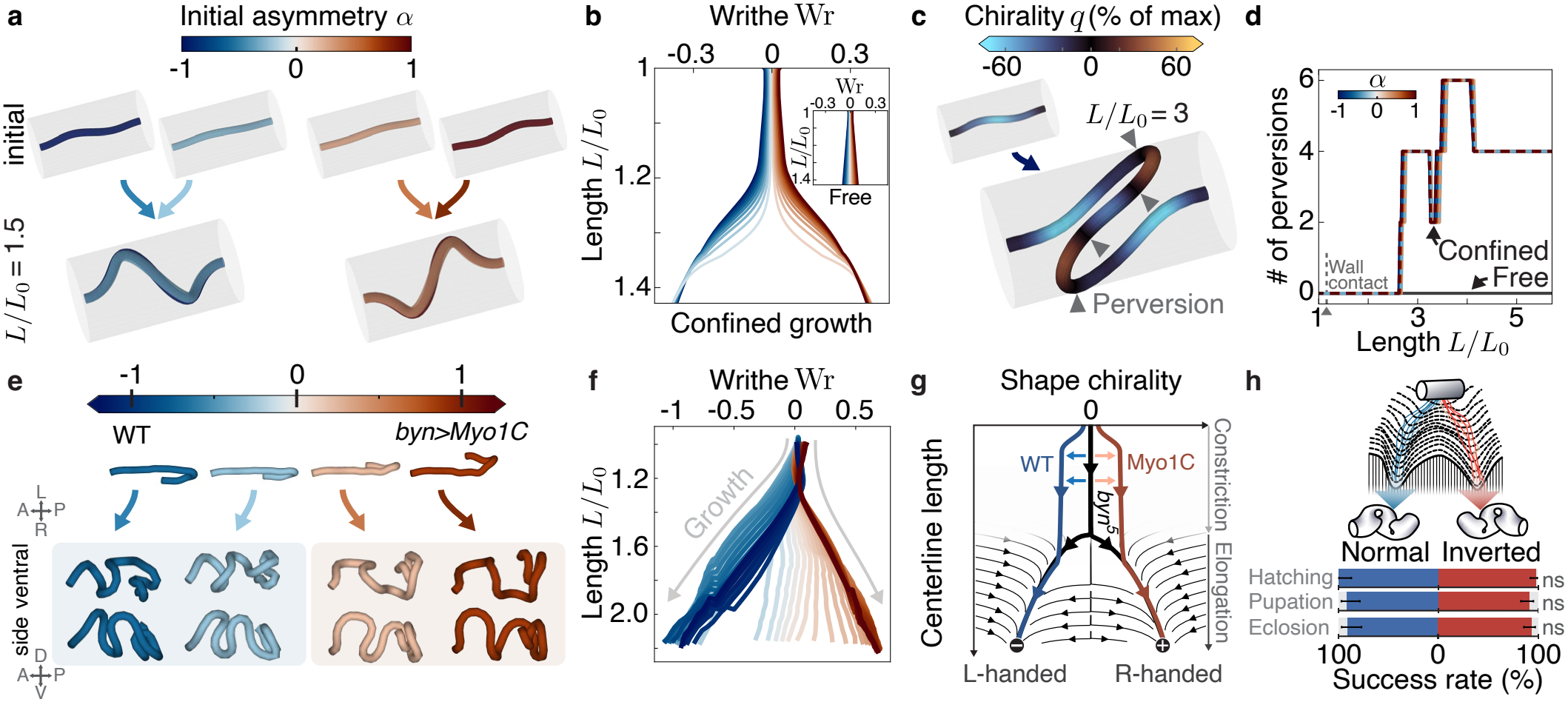
Buckling under confinement yields a mechanical route to canalization of developmental outcomes. **a**. Simulated confined, growing viscoelastic rods with fixed ends, started from shapes interpolating between left-handed (*α* = *−*1) and right-handed (*α* = +1), cluster into two opposite handednesses. **b**. Their Writhe Wr shows canalized dynamics during elongation, but only under confinement. Inset: growth in absence of confinement. **c**,**d**. Confined rods of initially uniform handedness become perverted as they grow, reducing torsion, as measured by the number of sign changes of the chirality; free rods do not. Examples in (**c**) for *α* = *−*1 at *L* = *L*_0_ and *L* = 3*L*_0_, colored by chirality. **e**,**f**. Midgut simulations from initial shapes interpolating between wild type (*α* = *−*1) and *byn>Myo1C* (*α* = +1) likewise cluster into two handednesses (**e**), with canalized Writhe dynamics (**f**). **g**. The constriction phase biases the initial condition; the elongation phase amplifies the asymmetry. **h**. Signaling and bistable mechanics together define a supracellular decision landscape with two canalized outcomes. Midguts of either handedness in the *Myo31DF* ^*L*152^ background pass developmental milestones at similar rates (hatching: *n* = 27 normal, *n* = 93 inverted; pupation and eclosion: *n* = 37 normal, *n* = 86 inverted). Error bars, Wilson 95% confidence intervals; *p*-values, two-sided Fisher’s exact test between phenotypes for each milestone; ns, *p >* 0.05.

The same holds for the midgut. Morphing the wild-type initial shape into the *byn>Myo1C* initial shape, and beyond (SI Sec. VII, Fig. S15; [80]), the simulated centerlines cluster into two groups of opposite handedness (Fig. 4e,f, Supp. Video 5), and the perversions shown in Fig. 1 follow from the mechanics of confined growth. Because asymmetric constrictions bias the initial condition toward one handedness, the cellular relay and elongation-driven buckling together act as an imperfect pitchfork bifurcation [81, 82]. This picture is consistent with the randomized handedness of *byn*^5^ and *HLH54F* ^Δ598^ midguts resulting from the loss of a biasing instruction (Fig. 4g). Other perturbations of the constriction stage, such as in *Antp* mutants, could produce additional morphological branches, which we leave to future work.

Finally, we test whether the stereotyped outcomes lead to differences in developmental fitness. We reared *Myo31DF*^*L152*^ embryos separated according to midgut handedness at stage 16. Both chiralities arise in this null background, avoiding potential confounding effects from variability in GAL4/UAS expression or timing. Both embryo cohorts–those with normal coiling and inverted coiling–pass developmental milestones of hatching, pupation, and eclosion at statistically indistinguishable rates under laboratory conditions (Fig. 4h). The mechanics of elongation under confinement thus funnels high-dimensional shape dynamics into two canalized out-comes, providing an explicit mechanism for the bistable canalization of organ geometry (Fig. 4h).

## DISCUSSION

Geometric constraints are a key tool for engineering complex shapes, from soft robots [83, 84] to stem-cell-derived tissues [85, 86]. Our work shows how, during development, the combination of long-range inter-organ cellular communication and supracellular mechanical interactions can similarly enable the emergence of stable 3D organ form. In the midgut, a weak chiral cue relayed between organs sets the direction of a buckling instability, and confinement funnels the growing organ into one of two stable forms. Such mechanical instabilities provide positive feedback and bistability with-out dedicated genetic regulation, in animals, plants and bacteria alike [25, 87–90]. Long-range mechanical coupling lets local control produce global shape changes while remaining robust to cell-scale heterogeneity and noise [22, 25, 91, 92]. Separating the selection of an outcome, by molecular cues, from its amplification and stabilization by global mechanics, may thus be a general design principle for building reproducible multicellular form.

Because mechanics alone permits two stable outcomes, a cue is needed to select one, which raises the question of whether handedness matters for function. Elongation increases the gut’s absorptive surface area and transit time [93], and accommodating this length within the body requires coiling, much like packing a rope into a confined space [94] or a viral genome into its capsid [95]; yet this geometric constraint does not favor either handedness. Consistent with this, normal and inverted midguts develop to adulthood at indistinguishable rates (Fig. 4h), suggesting that neither handedness may be functionally superior. Reliable handedness may instead matter for reproducible packing alongside other organs, and for avoiding pathological configurations such as volvulus, the twisting of the gut on itself in vertebrates [96].

Beyond the mechanics of coiling, our results support a model in which a multilayer cellular relay transmits a chiral bias from the hindgut epithelium to the midgut muscles via migrating myoblasts. While induction of left–right asymmetry from a spatially distinct organizer has several precedents [38, 97–100] the induction here appears to occur through a migratory cell population, raising several questions. Myoblasts may inherit a chiral polarity from interactions with the hindgut epithelium, or alternatively respond to an asymmetric signal emitted there. How the chiral bias emerges from Myo1C in the hindgut cells and how LVM cells then bias the direction of constrictions is also unclear: the left and right sides of the midgut deform differently (Fig. 2e), yet longitudinal muscle precursors on the two sides appear morphologically similar (Fig. S1), leaving open whether this asymmetry stems from differences in circular muscle contractility or in tissue material properties. Finally, our perturbations target Myo1C, and whether endogenous Myo1D sets wild-type handedness through the same relay remains to be tested [42].

Altogether, our results trace a program for organ shape distributed across tissues and scales, in which both cellular migration and tissue mechanics enable long-range propagation of molecularly-encoded chirality to 3D organ form. Traditionally, phenotypic switching has been understood through Waddington’s epigenetic landscape in a high-dimensional space of gene expression states [15, 16]. Augmenting this cellular picture with tissue mechanics allows us to understand the collective decision-making of multilayer tissues. It is fitting that Waddington’s metaphor, which originally appealed to mechanical intuition by picturing distinct phenotypes as marbles rolling down hills and valleys, here describes the mechanics of organ shape itself.

## ACKNOWLEDGMENTS

Sofia Villarreal conducted pilot Myo1C overexpression experiments. We are grateful for discussions with Sally Horne-Badovinac, Lucy O’Brien, Boris Shraiman, Suraj Shankar, Ousmane Kodio, Eric Wieschaus, Ed Munro, and the Cummings 9th-floor fly community. Research reported in this publication was primarily supported by the Eunice Kennedy Shriver National Institute Of Child Health & Human Development of the National Institutes of Health under Award Number R00HD110675. This research was also supported in part by the Physics Frontier Center for Living Systems funded by the National Science Foundation (PHY-2317138) and by grants from the NSF (DMS-2235451) and Simons Foundation (MP-TMPS-00005320) to the National Institute for Theory & Mathematics in Biology. FB thanks the Max Planck Society for funding. FB and NR acknowledge participation in the KITP workshop “Geometry and intercellular interactions in morphogenesis,” supported by the NSF PHY-2309135 and Gordon and Betty Moore Foundation Grant No. 2919.02. NR acknowledges support from the University of Chicago’s Biological Physics Fellowship and Center for Living Systems Fellowship. AF acknowledges support from the University of Chicago’s Bloomenthal Fellowship and the National Science Foundation Graduate Research Fellowship Program under Grant No. 2140001. Computing resources were provided by the University of Chicago Research Computing Center. Claude (An-thropic) was used for software assistance and copyediting in the later stages of the project.

## DATA AVAILABILITY STATEMENT

Data supporting the results in this article will be made available at Figshare.

## CODE AVAILABILITY STATEMENT

Code is available at https://github.com/MitchellLabCode/midgutChirality.

## METHODS

### 1. *Drosophila* husbandry and genetics

All *Drosophila melanogaster* stocks and crosses were kept at 25 ^*°*^C or 22 ^*°*^C. For imaging, embryos were laid on apple juice-agar plates smeared with fresh yeast paste. *w* ^1118^ was used as the wild-type strain. Light-sheet samples were held at 22 ^*°*^C during acquisition, and confocal imaging was performed at room temperature.

Transgenic stocks used in this study were: *UAS-Myo1C-RFP* (II) and *UAS-Myo1C* (III) (gifts from Stéphane Noselli; [42]); *Myo31DF*^*L*152^ (II) (gift from Kenji Matsuno; [37]); *byn-GAL4, klar* (III) (derived from *byn-GAL4* /*TM3, ftz-lacZ*, gift from Sebastian Streichan); *klar* (III) (gift from Eric Wieschaus; [101]); *Hand-GFP* (X) (gift from Zhe Han; [102]); *miple2-GAL4* (III) (gift from Ruth Palmer; [103]; [104]); *lab-GAL4* (III) and *Ubx-GAL4* (III) (gifts from Ben White and Ellie Heckscher; [105]); *NP5021-GAL4* (X) (DGRC 113545)). The following stocks were obtained from Bloomington *Drosophila* Stock Center: *byn*^5^ (III) (26186); *Df(2R)HLH54F*^Δ598^ (II) (91550); *HLH54Fd-GAL4* (II) (91633); *HLH54F*.*LVM-RFP* (II) (92001); *UAS-hid*.*Z, UAS-Stinger* (II) (65408); *48Y-GAL4* (II) (4935); *Mef2-GAL4* (II) (63842); nanos*-GAL4* (II) (4442); *wg-GAL4* (II) (83627); *Antp-GAL4* (X) (26817); *H2Av:iRFP* (II) (94935); *UAS-jGCaMP8s* (II) (92593); *Mef2-GAL4, UAS-mCD8*.*mRFP* (II) (26882); *UAS-mCherry*.*CAAX* (59021); *UAS-Lam*.*GFP* (III) (7378); *H2Av:iRFP* (94936; [106]); *20XUAS-IVS-jGCaMP8s* (III) (92593); *UAS-SERCA*.*R751Q*.*tdTomato* (II) (58972). Full experimental genotypes are listed in Table S1.

### 2. Live analysis of embryo handedness

For live analysis of embryos in the Myo1C overex-pression screen, the *byn*^5^ and *HLH54F* perturbations (Fig. 3), and the comparative developmental fitness assay (Fig. 4d), embryos were either submerged in Halocarbon 27 oil (Sigma-Aldrich) or dechorionated in bleach and mounted in water on a 35 mm glass-bottom dish (Cellvis). Midgut and hindgut handedness was scored by imaging embryos on Zeiss AxioZoom.V16 or Cephla SQUID microscopes. Blue excitation (470/40 nm bandpass for the AxioZoom.V16, 488 nm laser for the Cephla) was used to detect yolk autofluorescence and to identify embryos lacking fluorescent balancer chromosomes (*CyO, dfd-GMR-nvYFP*; *CyO, GAL4-Kr*.*C, UAS-GFP*.*S65T*; and *TM6B, Dfd-eYFP*), and green excitation (587/25 nm band-pass for the AxioZoom.V16, 568 nm laser for the Cephla) was used to detect UAS-Myo1C-RFP. Midgut chirality was scored at stage 16 as normal, inverted or abnormal in embryos that successfully formed all three midgut constrictions. Chirality was scored as abnormal when connections between midgut chambers were representative of both chiralities. Error bars represent 95% confidence intervals calculated using the Wilson score method, and *p*-values were calculated using Fisher’s exact test to assess differences in inversion frequency.

### 3. Immunofluorescence and tissue staining

Mouse anti-Fas3 (7G10, 1:50) and mouse anti-Cut (2B10, 1:10) antibodies were obtained from the Developmental Studies Hybridoma Bank. anti-Fas3 stains were used to visualize visceral mesoderm (Fig. 1c), and anti-Cut stains were used to score reduced sizes of malpighian tubules and hindguts in homozygous *byn*^5^ embryos (Fig. 3b). Embryos were collected on apple juice–agar caps, transferred to mesh baskets with a damp brush, dechorionated in 50% bleach for 2 min and rinsed in water. Dechorionated embryos were fixed for 15 min at room temperature with rocking in a two-phase mixture of heptane and 4% paraformaldehyde in PBST (1*×* PBS, 0.1% Triton X-100). After removal of the aqueous phase, embryos were devitellinized by adding an equal volume of methanol and vortexing; devitellinized embryos were washed three times in methanol and stored in methanol at *−*20 ^*°*^C until staining. For immunostaining, embryos were rehydrated with two 15-min washes in PBST and blocked for 1h at room temperature in blocking buffer (1% BSA in PBST). Then, embryos were incubated with primary antibodies, which were diluted in 1:1 blocking buffer:PBST, overnight at 4^*°*^C with rocking, then washed four times in PBST and incubated with secondary antibodies in 1:1 blocking buffer:PBST between 1 and 2 hours at room temperature. All subsequent steps were performed protected from light. Embryos were washed three times in PTW (1 *×*PBS, 0.1% Tween-20). Stained embryos were optically cleared in increasing concentrations of glycerol up to 90%, then mounted and imaged in glass-bottom dishes under Vectashield^®^ with DAPI.

### 4. Confocal and widefield imaging

Confocal imaging was done on inverted Zeiss LSM 980, Zeiss LSM 880, and Leica SP8 laser scanning confocal microscopes, using solid-state and diode laser lines. Wide-field imaging was done on upright Zeiss Axio Zoom V.16 microscopes, using brightfield and widefield epifluorescence. All embryos were staged before or during stage 15, prior to the onset of coiling, unless otherwise noted.

### 5. Light-sheet imaging and processing

Volumetric light-sheet imaging was performed on a Bruker Luxendo MuVi (Multi-View) SPIM light-sheet microscope with 488 nm, 561 nm, and 642 nm laser lines to excite the following: Hand-GFP (circular visceral muscle nuclei) and Dfd-eYFP; mCherry.CAAX, HLH54F.LVM-RFP (longitudinal visceral muscles), and Myo1C-RFP; and H2Av:iRFP (endodermal nuclei), respectively. Embryos were mounted in 1.2% agarose containing 0.5 *μ*m fluorescent microspheres (TetraSpeck, Invitrogen). Six views, at 60^*°*^ increments, were acquired at every time point. Views were then registered, fused and deconvolved using the Multiview Registration and Deconvolution pipeline [107], with the fluorescent microspheres as interest points for registration. The final deconvolution step adapts the iterative Richardson-Lucy algorithm to multiview datasets [108], using a measured point-spread function.

We then performed pixel classification on the fused, deconvolved volumes using the `ilastik` “Autocontext (2-stage)” workflow, trained on manual annotations, to identify the basal surface of the endoderm [109]. A level-set method then obtained a 3D segmentation of this surface, which we converted to triangulated meshes using methods in [53].

### 6. Surface segmentation and parametrization

Meshes of the basal side of the endodermal surface of the midgut were parametrized using `TubULAR` [53], which establishes longitudinal and circumferential coordinates across time and provides mappings between these surface coordinates and their embedding in 3D. Anterior and posterior endcaps were removed to transform each closed surface into a topological cylinder, and a longitudinal cut path joining the two ends was defined automatically along the dorsal side of the tissue. We generated stabilized, quasiconformal pullbacks that encode the longitudinal and circumferential parameterization [53]. We used these pullbacks to construct an orientation frame from the pattern of circular visceral muscle nuclei along the organ, shown in yellow in Fig. 1d. The construction of this frame from images is detailed in SI Sec. IV.

### 7. Viscoelastic rod model

We use a Cosserat rod model to simulate the evolution of the midgut in the coiling phase, during which we approximate the organ as an elongating, unshearable one-dimensional rod that resists bending and twisting, and whose stress-free rest configuration adapts to the gut’s current state at a set remodeling rate. We label points along the gut by their fractional arclength 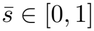, and describe the rest configuration by its length density 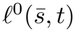, such that the rest arclength *s* satisfies 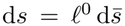, and by its curvature 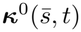. In this section, primes denote derivatives with respect to the rest arclength, 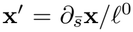, so that **x**^*′*^ = 1 when the gut is at its rest length. Deviations from the rest configuration generate internal forces **n** = *Q*^*⊤*^**S**(*Q***x**^*′*^*−* **e**^3^) and torques **m** = *Q*^*⊤*^**B***Q*(***κ****−* ***κ***^0^), where the rows of *Q* are the frame vectors **d**_*i*_, **e**^3^ = (0, 0, 1)^*⊤*^, and the Darboux curvature vector is defined by 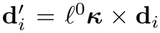 [110]. The matrices **S** and **B** = diag(*B*_1_, *B*_1_, *B*_3_) are diagonal shear/stretch and bending/twist stiffness matrices, with *B*_1_ the bending stiffness and *B*_3_ the twisting stiffness. Together with external forces **f** ^ext^ due to self-contact and collisions with the anatomical cavity, these drive the motion of the centerline and the rotation of the frame through the over-damped balance equations

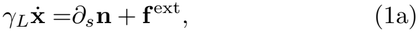

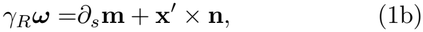

with translational and rotational drag coefficients *γ*_*L*_ and *γ*_*R*_, and the frame evolving as *∂*_*t*_**d**_*i*_ = ***ω****×* **d**_*i*_. To account for the observed elongation of the centerline and for remodeling of the rest configuration, the rest configuration evolves according to

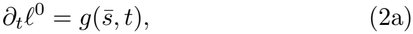

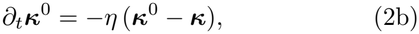

where *η* is the remodeling rate and the growth field 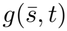 is measured from the elongation of the experimental centerlines. The regime most faithful to the experiments combines fast remodeling with relatively fast growth, such that the midgut behaves in a fluid-like manner, with *ητ*_*E*_ = 10 and *T/τ*_*E*_ = 3, where *T* is the duration of the simulation and 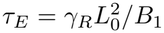 the elastic relaxation time of a midgut of initial length *L*_0_ (Fig. S13). Simulations are implemented in `julia` and account for contact with the anatomical cavity surrounding the midgut, self-contact between gut segments, and experimentallyderived growth rates and boundary conditions. More theory and implementation details are provided in SI Sec. VI.

## Notes

### Competing Interest Statement

The authors have declared no competing interest.

