## Supplementary Information for "Mechanical canalization of chiral gut morphogenesis"

#### CONTENTS


I. Experimental genotypes 2

II. Features of $byn^5$ mutant embryos 2

III. Organ surface reconstruction 3
1. `ilastik` pixel classification 3
2. `TubULAR`-based surface parameterization 4
3. Time alignment 5
4. Surface kinematics 6

IV. Framed curves 7
1. Geometric preliminaries 7
2. Curve construction 8
3. Construction of material frame 8
4. Measures of deformation 9

V. Measuring centerline chirality and topology 10
1. Local measure 10
2. Quasi-local measure 12
3. Link, Writhe, Twist and topological constraint 12

VI. Viscoelastic Cosserat rod dynamics 14
1. Continuum Cosserat rod theory 14
2. Discretization of the Cosserat rod 15
3. Growth-rate field from data 16
4. Boundary conditions 17
5. Remeshing 17
6. Self-contact 17
7. Anatomical confinement 18
8. Parameter sweeps 20
9. Effect of confinement and boundary orientation 22

VII. Canalization of midgut chirality in interpolated shapes 22
1. Initial condition interpolation and canalization 22
2. Interpolation of growth rates 24
3. Interpolation of self-contact and confinement 25

VIII. Scaling argument for confinement-driven chirality selection 25

IX. Supplementary Video captions 27

References 28

### I. EXPERIMENTAL GENOTYPES

TABLE S 1. Experimental genotypes used in this study

| Experiment | Figure | Experimental genotype | Parental cross |
|---|---|---|---|
| Representative image of normal coiling | Fig. 1c | $w^{1118}$;+;+ | $w^{1118}$;+;+ |
| Representative image of inverted coiling | Fig. 1c | *w*; +; *bynGAL4*, *UAS-Myo1C/+* | Female: *w*;+; *bynGAL4*, *UAS-Myo1C/TM6B, PDfd-eYFP* Male: $w^{1118}$;+;+ |
| LVM visualization | Fig. 1b | *y*, *w*; *HLH54F.LVM-RFP* | Female/Male: *y*, *w*; *HLH54F.LVM-RFP* |
| $byn^5$ — Live scoring | Figs. 3b & S2 | +;+;$byn^5$, *e* | Female/Male: +;+;$byn^5$, *e/ TM6B, PDfd-eYFP* |
| $byn^5$ — Light-sheet | Fig. S2 | *w*, *Hand-GFP/w*; *HLH54F.LVM-RFP/H2Av:iRFP*; $byn^5$, *e*, *klar* | Female: *w*; *HLH54F.LVM-RFP*; $byn^5$, *e*, *klar/TM3, PActGFP, Ser, klar* Male: *w*, *Hand-GFP*/Y; *H2Av:iRFP*; $byn^5$, *e/ TM6B, PDfd-eYFP* |
| LVM null | Fig. 3c | *w*; $Df(2R)HLH54F^{\Delta 598}$ | Female/Male *w*; $Df(2R)HLH54F^{\Delta 598}$*/CyO, KrGAL4, UAS-GFP.S65T*; + |
| LVM null with Myo1C overexpression | Fig. 3c | *w*; $Df(2R)HLH54F^{\Delta 598}$; *bynGAL4*, *UAS-Myo1C/+* | Female *w*; $Df(2R)HLH54F^{\Delta 598}$*/CyO, KrGAL4, UAS-GFP.S65T* Male: *w*; $Df(2R)HLH54F^{\Delta 598}$/CyO, dfd-GMR-nvYFP; *bynGAL4*, *UAS-Myo1C/TM6B, PDfd-eYFP* |
| LVM ablation (*HLH54F*>hid, Stinger) | Fig. 3c | *w*, *Hand-GFP/w*; *UAS-Stinger*, *UAS-hid.Z/HLH54Fd-GAL4*; *klar/H2Av:iRFP* | Female: *w*, *Hand-GFP*; *HLH54Fd-GAL4*/CyO, dfd-GMR-nvYFP; *klar* Male: *w*; *UAS-Stinger*, *UAS-hid*/CyO, dfd-GMR-nvYFP; *H2Av:iRFP* |
| Light-sheet live — WT | Figs. 1 & 2 | *w*, *Hand-GFP/w*; *UAS-mCherry.CAAX/+*; *byn-GAL4*, *klar/H2Av:iRFP* | Female: *w*, *Hand-GFP*;+; *bynGAL4*, *klar/TM3, Sb, klar* Male: *w*; *UAS-mCherry.CAAX.S*; *H2Av:iRFP* |
| Light-sheet live — Myo1C overexpression | Figs. 1 & 2 | *w*, *Hand-GFP/w*; *UAS-Myo1C-RFP/+*; *byn-GAL4*, *klar/H2Av:iRFP* | Female: *w*, *Hand-GFP*;+; *bynGAL4*, *klar/TM3, Sb, klar* Male *w*; *UAS-Myo1C-RFP/CyO*; *H2Av:iRFP* |
| Comparative developmental fitness | Fig. 4h | *w*; $Myo31DF^{L152}$;+ | Female/Male *w*; $Myo31DF^{L152}$;+ |
| Calcium imaging | Fig. 2a | *w*; *+/CyO, Mef2-GAL4, UAS-mCD8.mRFP*; *20XUAS-IVS-jGCaMP8s/+* | Female: *w*; $wg^{Sp-1}$*/CyO, Mef2-GAL4, UAS-mCD8.mRFP*; *20XUAS-IVS-jGCaMP8s* Male: $w^{1118}$;+;+ |
| *byn-GAL4* expression domain | Fig. S1 | *w*; *UAS-mCherry.CAAX.s/+*; *UAS-Lam.GFP/bynGAL4*, *klar* | Female: *w*; +; *bynGAL4*, *klar / TM3, Sb, klar* Male: *w*; *UAS-mCherry.CAAX.s*; *UAS-Lam.GFP* |

### II. FEATURES OF $BYN^5$ MUTANT EMBRYOS

To better understand the presence of residual longitudinal visceral muscle (LVM) structure in $byn^5$ mutants, we performed light-sheet microscopy on $n = 10$ individuals during stage 16 or 17 with fluorescent proteins marking each of the midgut's three tissue layers: H2Av:iRFP, Hand-GFP, and HLH54F.LVM-RFP fluoresce in endodermal nuclei, CVM nuclei, and LVM nuclei, respectively. The deconvolved microscopy datasets were trained on in `ilastik`, processed in `TubULAR`, and smoothed in `MeshLab` to construct three-dimensional (3D) textured meshes of the midgut surfaces (see Sec. III 1 for details of this workflow). These meshes revealed that the coverage of residual LVM cells is highly variable in the homozygous mutants, yet less extensive than that of heterozygous control siblings (Fig. S2a-d). Further imaging of $byn^5$ embryos from stage 14 until the onset of coiling in stage 16 showed that the majority of midguts have missing or abnormal constrictions (Fig. S2e).

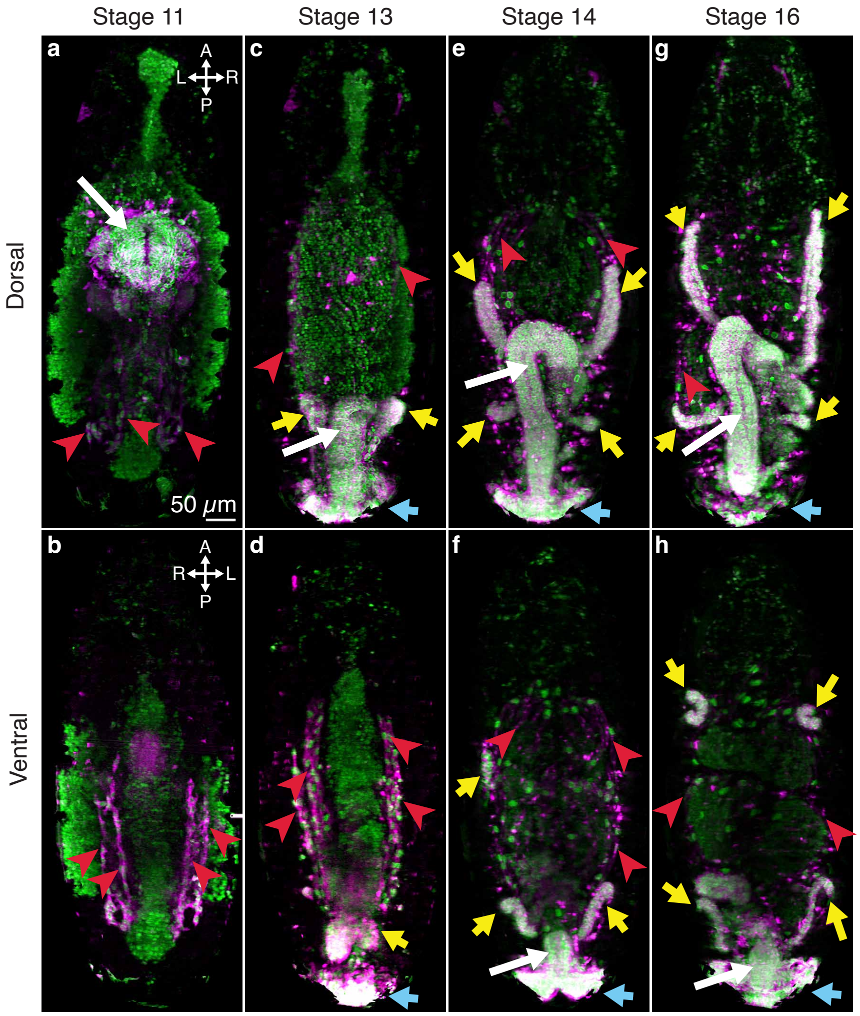


FIG. S1. *byn-GAL4* domain includes hindgut (white arrows) and longitudinal muscles (red arrowheads), along with malpighian tubules (yellow arrows) and anal pads (cyan arrows), as shown in deconvolved multi-view light-sheet microscopy data of a *byn>mCherry.CAAX,Lam.GFP* embryo from stages 11 through 16. Images are maximum intensity projections of half of the embryo volume, from the middle of the embryo to either the dorsal (top) or ventral (bottom) side. **a-b**. At stage 11, UAS-driven fluorophore expression is visible in the hindgut primordium and longitudinal visceral muscle precursors. **c-d**. Expression at stage 13 includes the elongating hindgut, longitudinal muscles, Malpighian tubules, and anal pad. **e-f**. Fluorophore expression at stage 14 is detected in the same tissues. **g-h**. Expression at stage 16 is similar to stage 14. Asymmetric midgut coiling is visible in the ventral view.

#### III. ORGAN SURFACE RECONSTRUCTION

##### 1. `ilastik` pixel classification

After multi-view deconvolution [5], we use `ilastik`'s Two-stage Autocontext workflow [1] to classify pixels before volumetric segmentation. In the first stage, we separately labeled endoderm, yolk, visceral mesoderm and surface laying just outside the basal surface of the endoderm, and other tissues. As shown in Fig. S3b, the second stage uses four labels: one for the endoderm and enclosed yolk, one for musculature and background, and one each for anterior and posterior endcaps connecting the midgut to the proventriculus and pylorus, respectively. The endcaps close the gut at its anterior and posterior boundaries, enabling subsequent segmentation via level-sets segmentation. The first channel's output probability maps therefore specify the likelihood that a given pixel is inside or outside the gut, proximal to the visceral muscle layer. These probabilites are then input into a `TubULAR` workflow that segments the volume by minimizing a Chan–Vese functional [6]. Marching cubes converts this segmentation into a triangulated mesh. These meshes are then smoothed using a series of `MeshLab` filters. Namely, we perform multiple rounds of HC

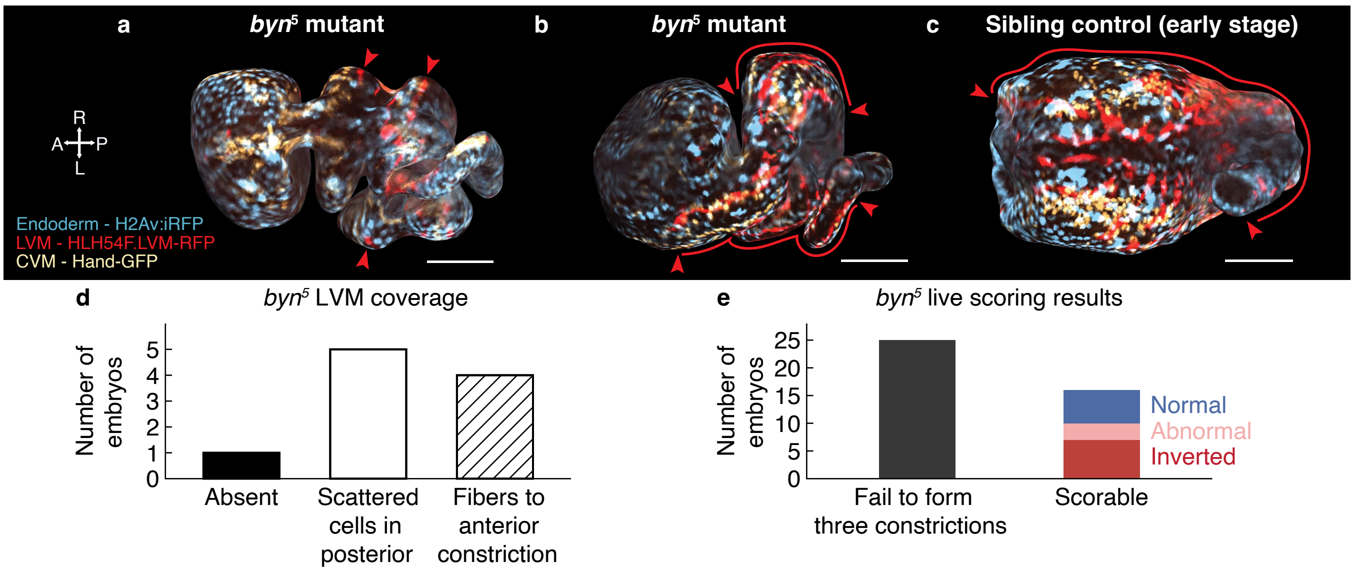


FIG. S2. *byn*$^5$ mutant embryos have reduced LVM coverage and often lack at least one midgut constriction. **a-b-c**. Dorsal views of 3D textured surfaces of *Hand-GFP*;*HLH54F.LVM-RFP*/*H2Av:iRFP* embryos imaged by light-sheet microscopy, scale bars 50 $\mu$m. **a**. Stage 16 *byn*$^5$ embryo with abnormal constrictions and LVM cells (red arrowheads) scattered throughout the posterior midgut. **b**. Stage 16 *byn*$^5$ embryo with multiple LVM fibers (red arrowheads) spanning from the posterior to the anterior-most constriction. **c**. In a Stage 15 control sibling heterozygous for *byn*$^5$, LVM fibers (red arrowheads) extend through the full anterior portion of the midgut. **d**. LVM coverage scored in *byn*$^5$ embryos imaged via light-sheet microscopy ($n = 10$). LVM coverage is variable but is not observed to reach as far as control siblings or wild-type embryos. **e**. Live analysis of homozygous *byn*$^5$ embryos imaged from stage 14 until midgut coiling ($n = 41$), scored by ability to form three constrictions. Embryos that failed to form three constrictions were more numerous but not scored for handedness, while constriction-forming embryos were included and scored in Main Fig. 3b.

Laplacian smoothing to reduce local surface irregularities while preserving the overall geometry. Poisson-disk sampling then provides a resampled set of surface points, followed by screened Poisson surface reconstruction to generate a smooth, continuous mesh. 10 iterations of LS3 Loop subdivision refine the reconstructed surface.

##### 2. `TubULAR`-based surface parameterization

We use `TubULAR` [3] to define a surface parametrization $\mathbf{X}(u, \phi)$ of each midgut over time. Meshes of the basal endoderm midgut surface (the outermost gut surface layer laying within the visceral muscle) were parametrized to establish longitudinal and circumferential coordinates across time and provide mappings between these surface coordinates and positions in 3D space.

Anatomical landmarks were identified from separately trained `ilastik` probability maps. First, we labeled anterior, posterior, and dorsal landmarks to align meshes into a global anatomical frame (APDV) coordinate system. We identified the anterior connection with the proventriculus, a point on the forming dorsal vessel, and a point posterior to the midgut at a reference time point at the beginning of stage 15. All surface meshes were subsequently aligned within this common APDV frame. We then identified the midgut's connection with the proventriculus and its connection with the pylorus at each timepoint to define endpoints of the gut.

For each timepoint we extracted a coarse centerline connecting these endpoints by fast marching through the 3D distance transform of the segmentation. This centerline serves only to constrain the subsequent surface cut and is not used for geometric measurement. To transform each closed surface into a topological cylinder, we removed the largest connected component of the mesh lying within 19.5 $\mu$m of each endpoint, measured as geodesic distance along the surface. `TubULAR` was then used to generate a cut along the long axis of the mesh and defined corresponding longitudinal–circumferential coordinates for each time point via conformal and stabilized quasiconformal maps.

In addition, we use these pullbacks to define a frame constructed by the circular visceral muscles patterned along the length of the organ, as shown in yellow in Fig. S3d and Main Fig. 1. The construction of this frame from images is detailed in Sec. IV.

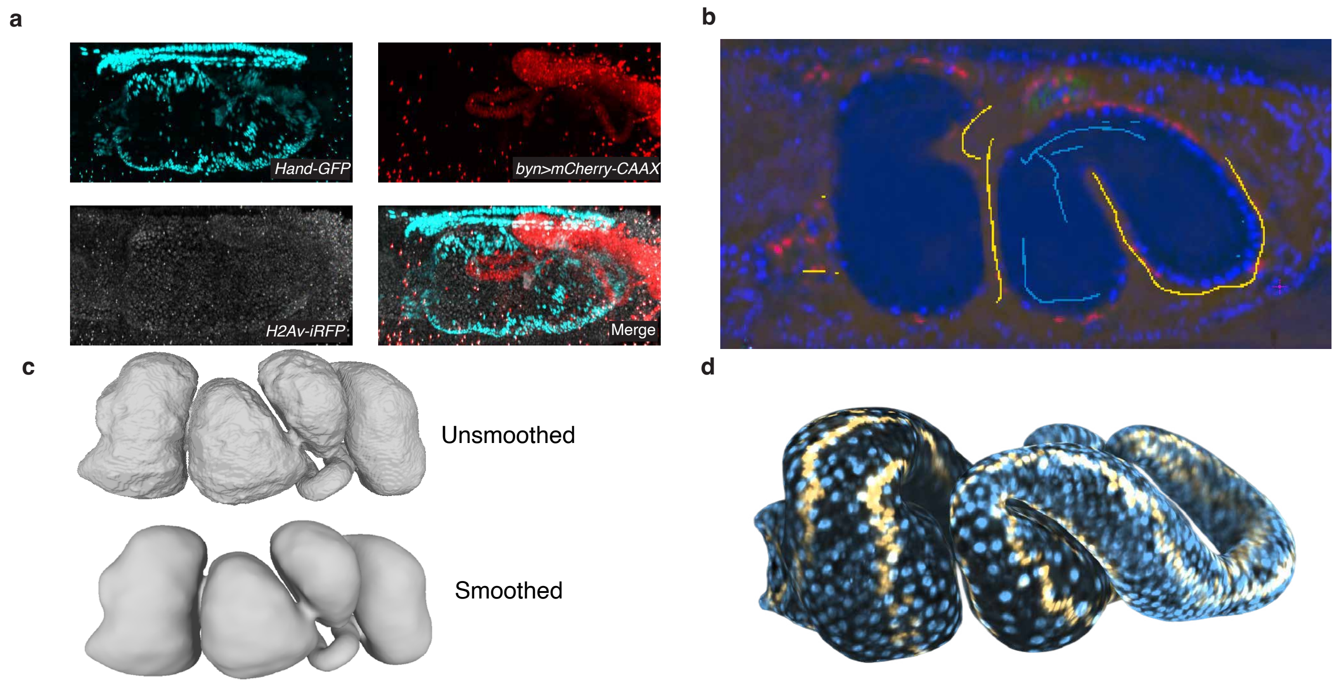


FIG. S3. Reconstruction of the *Drosophila* midgut surface as a textured triangular mesh from multicolor, multiview light-sheet images. **a**. Raw data for kinematic calculations in this paper were taken using the Bruker/Luxendo MuVi SPIM Multi-view light-sheet microscope, and then deconvolved. **b**. We use deconvolved, downsampled volumetric data as inputs to random-forest-based pixel classifier `ilastik` [1] in its two-stage autocontext feature to extract the basal surface of the *Drosophila* midgut. **c**. The probability matrix extracted from `ilastik` then allows us to obtain 3D representations of the shape, which we then smooth using a series of filters in Meshlab [2]. **d**. To quantitatively study cellular surface dynamics, we finally map the 3D image to a surface texture using `TubULAR` [3].

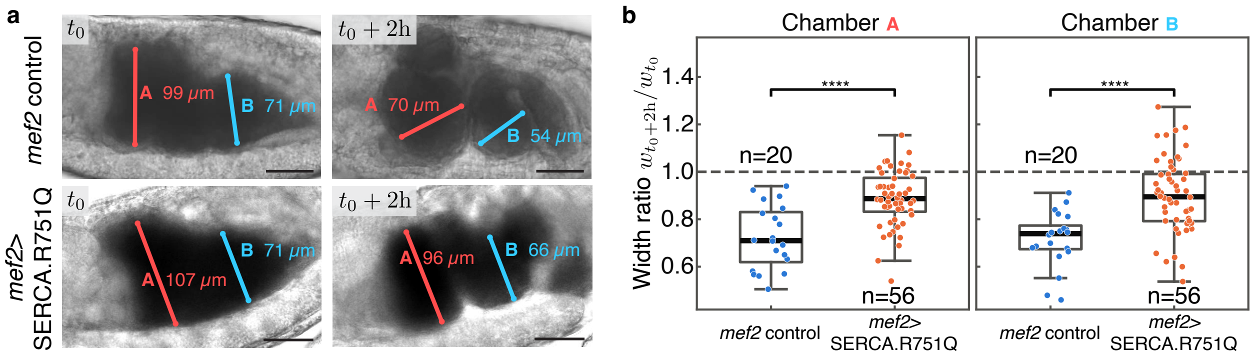


FIG. S4. Calcium activity drives decrease in radius. **a.** Reanalysis of calcium pump inhibition data by heat-dependent calcium pump inhibition data using modified *UAS-SERCA.R751Q* dominant neomorphic perturbation from Ref. [4]. We measure and compare the width $w_t$ of anterior and posterior chambers (A and B) at time $t = t_0$ and $t_0 + 2$h. **b.** Perturbation of calcium activity leads to a significant perturbation in the ability of muscles to contract the chamber radii. $p$-values computed using the two-sided Mann-Whitney U test on the ratios. Chamber A, $p = 3.4 \times 10^{-5}$. Chamber B, $p = 7.6 \times 10^{-6}$.

##### 3. Time alignment

To place independently imaged embryos on a common developmental timeline, we align datasets by the temporal progression of the midguts' radius profiles. We construct the radius kymograph $R(s,t)$ for each dataset as follows. We extract the surface radius profile $R(s)$, which is the $\phi$-averaged distance to the mean position of the embedding coordinates of a curve of constant $s$. We find the three constriction (fold) positions using `TubULAR`'s fold-detector (now packaged in the updated GitHub release), by identifying stable local minima in the radii. These positions define

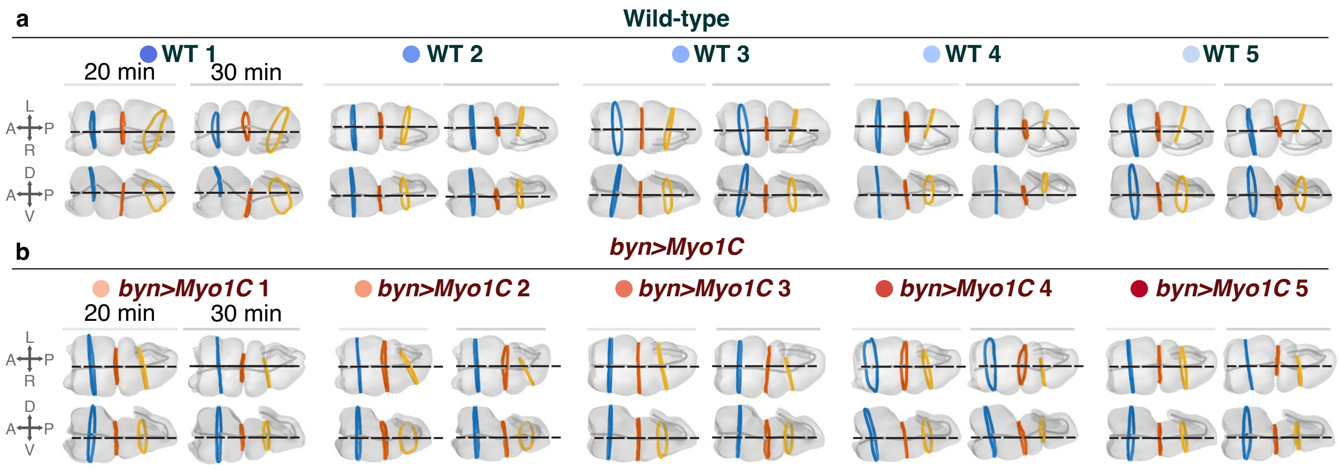


FIG. S5. Constrictions break symmetry. Centerlines and constrictions at $t = 20$ and $t = 30$ min in the reference timeline for wild-type (**a**) and *byn>Myo1C* samples (**b**). $t = 0$ min marks the onset of first constriction in sample WT 2.

chamber boundaries along the normalized anteroposterior coordinate $s/L \in [0, 1]$ at each timepoint. This gives us a radius kymograph $R(s, t)$ for each dataset.

To make profiles comparable across embryos of differing size and fold placement, each kymograph is remapped onto a fixed fold-anchored grid $\tilde{s}$: within each of the four chambers we re-bin the data to a prescribed number of points ($n = 100$), so that the folds always land at the same position $\tilde{s}$ regardless of their absolute position. We then group datasets by genotype (wild-type and *byn>Myo1C*) and, within each group, align every embryo to a designated reference embryo by cross-correlating the fold-normalized radius kymographs: for each candidate relative time shift, we slide one kymograph against the reference, compute a Pearson correlation between the two embryos' radius time-courses at each spatial position over the overlapping time window, and average these across all spatial positions to score that shift. To make this correlation measure sensitive to the shape of the radius profile, rather than its scale, radius profiles are normalized by subtracting their mean and dividing by the standard deviation. We smooth the resulting correlation-versus-shift curve with a Savitzky–Golay filter and take the shift value at which the curve peaks as the optimal relative time shift for that embryo.

Finally, the two genotype references are aligned to one another using the same cross-correlation procedure, yielding a single cross-genotype offset. The resulting per-embryo shifts, relative to each genotype reference, and the cross-genotype offset are saved as the relative timing needed to register all datasets onto one timeline. We then use the onset of the first constriction in the reference WT sample to define the origin $t = 0$.

##### 4. Surface kinematics

We measure how the tissue surface deforms over time to query how reduction in midgut radius couples to elongation of its centerline. We tracked tissue motion by particle image velocimetry (PIV) on the stabilized material frame $(u, \phi)$ pullbacks described in Sec. III 2, using `PIVlab` [7, 8] through the `TubULAR` pipeline. The PIV is computed between consecutive frames ($\Delta t = 2$ min).

We then convert the variations in 3D displacements to velocities by taking the difference between displacement vectors between time $t$ and $t + \Delta t$,

$$\mathbf{v} = \frac{\mathbf{X}_{t+\Delta t}(s + \Delta s,\, \phi + \Delta\phi) - \mathbf{X}_t(s, \phi)}{\Delta t}, \tag{S1}$$

so that the 3D velocity $\mathbf{v}$ accounts for both motion within the surface (in-plane) and motion of the surface itself (out-of-plane). To reduce measurement noise, velocities were then temporally smoothed along Lagrangian pathlines using a triangular (tripulse) filter spanning five consecutive frames. Introducing the unit vector normal to the surface $\hat{\mathbf{n}}$, we can then decompose the velocity into its normal component $v_n = \mathbf{v} \cdot \hat{\mathbf{n}}$ and in-plane component $\mathbf{v}_\parallel = \mathbf{v} - v_n \hat{\mathbf{n}}$.

With this decomposition of the velocity, we can then estimate the local rate of change of a tissue patch of area $\delta A$ using the geometric relationship

$$\frac{1}{\delta A}\frac{\mathrm{d}(\delta A)}{\mathrm{d}t} = \tfrac{1}{2}\,\mathrm{Tr}\big[\mathbf{g}^{-1}\dot{\mathbf{g}}\big] = \nabla \cdot \mathbf{v}_\parallel - 2H v_n, \tag{S2}$$

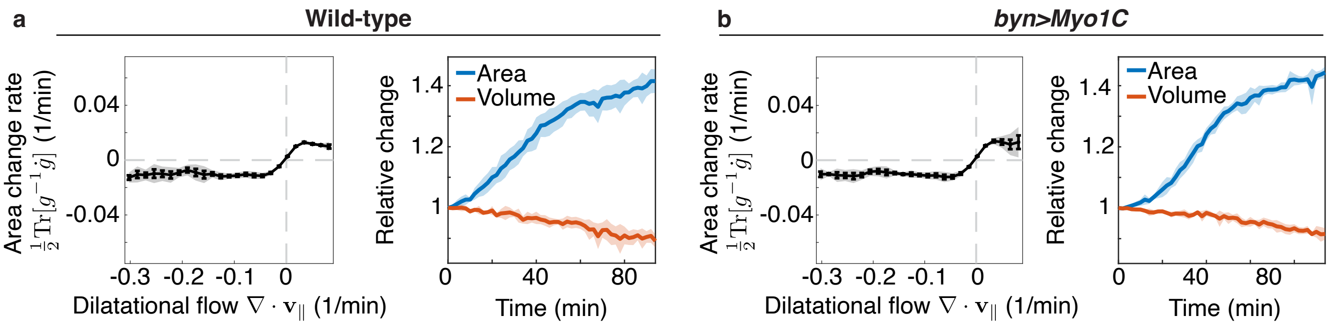


FIG. S6. Local area changes are modest compared to surface flows. **a.** Wild type, **b.** *byn>Myo1C*. Left: area change rate $\frac{1}{2}\mathrm{Tr}[g^{-1}\dot{g}]$ binned by the dilatational flow $\nabla\cdot\mathbf{v}_{\|}$. The area change rate's magnitude is bounded by 0.01 min$^{-1}$, an order of magnitude below the strongest flows ($|\nabla\cdot\mathbf{v}_{\|}|\approx 0.1$ min$^{-1}$). The sign follows that of the dilational flow. Because $\frac{1}{2}\mathrm{Tr}[g^{-1}\dot{g}] = \nabla\cdot\mathbf{v}_{\|} - 2Hv_n$, in-plane convergence is offset to within 10% by out-of-plane motion of the tissue ($2Hv_n$). Each data point reflects the instantaneous kinematics of local patches, each 1% of the organ length and 1% of the circumference, averaged across one quarter of the gut circumference, as in [3, 4]. Points show the mean in each bin; error bars indicate the standard error on the mean, shading standard deviation. Right: total surface area and enclosed volume of the midgut, relative to their values at $t = 0$. Over ~100 min the area grows by ~40% in both cases, while the volume falls by ~10%. $n = 5$ embryos per genotype. Time is aligned as indicated in Sec. III 3.

where $\mathbf{g}$ is the surface metric, $\dot{\mathbf{g}}$ its time-derivative, and $H = -\frac{1}{2}\nabla\cdot\hat{\mathbf{n}}$ the mean curvature. Consequently, a surface that conserves area locally must satisfy the relationship

$$\nabla\cdot\mathbf{v}_{\|} \approx 2Hv_n. \tag{S3}$$

We find that Eq. (S3) holds across both control (*byn>mCh-CAAX*) and *byn>Myo1C* midguts: its left- and right-hand sides are strongly correlated, with mean Pearson correlations of $\rho = 0.96 \pm 0.01$ and $0.97 \pm 0.02$ for control and *byn>Myo1C* midguts respectively, consistent with measurements focusing on earlier stages [4]. Beyond correlation, we find that the two sides of (S3) are nearly equal, with a total least squares slope of $0.97 \pm 0.01$ between the two quantities for both genotypes. The limited compressibility of the tissue surface provides a kinematic mechanism to convert circumferential contractility to elongation.

#### IV. FRAMED CURVES

To quantify the overall chiral shape of the embryonic midgut, we reduce the organ to the time-dependent space curves defined by its centerline $\mathbf{x}(s,t)$ and myocyte curves $\{\mathbf{x}_j(\lambda,t)\}_{j=1,2,3,4}$, with $\mathbf{x},\mathbf{x}_j \in \mathbb{R}^3$. In this section, we expose the construction of these curves and how we use these to represent the orientation of the organ surface by a rotation matrix $Q \in \mathrm{SO}(3)$. Together with the centerline, this defines a framed curve $(\mathbf{x},Q) \in \mathbb{R}^3 \times \mathrm{SO}(3)$. We then detail measures of deformation of these framed curves, before describing measure of chirality in Sec. V; the kinematic observations derived from these measurements form the basis for the viscoelastic Cosserat model of Sec. VI.

##### 1. Geometric preliminaries

Before detailing any construction from data, we introduce tools to manipulate the rotation matrices $Q$ that carry the orientation of rod segments. For a vector $\mathbf{v} \in \mathbb{R}^3$, we denote by $\mathrm{skew}(\mathbf{v})$ the skew-symmetric matrix representing the cross product, $\mathrm{skew}(\mathbf{v})\,\mathbf{w} = \mathbf{v}\times\mathbf{w}$ for all $\mathbf{w}\in\mathbb{R}^3$,

$$\mathrm{skew}(\mathbf{v}) = \begin{pmatrix} 0 & -v_3 & v_2 \\ v_3 & 0 & -v_1 \\ -v_2 & v_1 & 0 \end{pmatrix}, \tag{S4}$$

and by $[\cdot]_\times$ the inverse operation, $[\mathrm{skew}(\mathbf{v})]_\times = \mathbf{v}$, which reads $\mathbf{v} = (V_{32}, V_{13}, V_{21})$ from $V = \mathrm{skew}(\mathbf{v})$.

A $3\times 3$ rotation matrix $R$, with inverse $R^{-1} = R^\top$, can be represented by a vector $\mathbf{u}\in\mathbb{R}^3$ whose direction $\hat{\mathbf{u}} = \mathbf{u}/|\mathbf{u}|$ specifies the axis of rotation and whose norm $\theta = |\mathbf{u}|$ specifies the rotation angle. The two representations are related by the matrix exponential, $R = \exp(\text{skew}(\mathbf{u}))$, which can be computed with Rodrigues' rotation formula

$$R = I + \sin\theta\, U + (1-\cos\theta)\, U^2, \qquad U = \text{skew}(\hat{\mathbf{u}}). \tag{S5}$$

Conversely, the axis-angle representation $\mathbf{u}$ of a given rotation matrix $R$ is obtained from the matrix logarithm

$$\mathbf{u} = \log R = \begin{cases} 0 & \text{if } \theta = 0 \\ \frac{\theta}{2\sin\theta}[R - R^\top]_\times & \text{if } \theta \neq 0 \end{cases}, \quad \theta = \arccos\left(\frac{\text{tr}R - 1}{2}\right). \tag{S6}$$

These relations allow for efficient computation of curvatures and rotation rates.

#### 2. Curve construction

With these geometric preliminaries, we now detail the construction of the midgut centerline and the myocyte curves. At a fixed time $t$, given the surface parametrized as $\mathbf{X}(u,\phi) \in \mathbb{R}^3$, with $u \in [0,1]$ and $\phi \in [0,2\pi)$, we define the centerline at each position $u$ by the centroid of the section

$$\mathbf{x}(u) = \frac{1}{2\pi}\int_0^{2\pi} \mathrm{d}\phi\, \mathbf{X}(u,\phi). \tag{S7}$$

We use the arclength parametrization $s \in [0, L]$ rather than the surface longitudinal parameter $u$ by using the change of variable defined by the differential equation

$$\frac{\mathrm{d}s}{\mathrm{d}u} = |\partial_u \mathbf{x}(u)|, \quad s(0) = 0. \tag{S8}$$

To define the orientation of the midgut, we use the positions of the myocyte nuclei. These are physically coupled to the endodermal layer and can thus be used as fiducial markers for the material orientation of the midgut. The myocyte nuclei are arranged in four lines on the surface of the midgut. These are hand-drawn as parametrized curves $\lambda \mapsto (u(\lambda), \phi(\lambda))$ on the 2D projections of the 3D images onto the segmented surface mesh. The curves are then pulled back to 3D curves $\mathbf{x}_j(\lambda)$ (Fig. S7a-b).

All resulting curves (represented as polylines) then require additional smoothing to enable reliable estimation of curvatures. To smooth the data with prescribed low-pass filters in both space and time without inducing shrinkage, we use Taubin smoothing in space [9] and Gaussian smoothing in time.

#### 3. Construction of material frame

Once we have the centerline $\mathbf{x}$ and myocyte curves $\{\mathbf{x}_j\}_{j=1,2,3,4}$ at each time, we define the material frame of the curve by weighting the directions between centerline and myocytes through a dipolar-like vector field construction (Fig. S7c). In words, we find the intersection points $\mathbf{x}_j(s,t)$ of the myocyte curves with the sections $\phi \mapsto \mathbf{X}(s,\phi)$ and we assign 'charges' $c_j = \pm 1$ to these points, which generate an 'electric field' pointing from right to left (in the body axis orientation). The orientation of this field weights the relative importance of the myocytes through their inverse-square distance to the centerline. Formally, we define the local orientation through the vector

$$\mathbf{M}(s,t) = \sum_{s'=s-n_a}^{s+n_a} \sum_{j=1}^{4} q_j\, P^\perp_{\mathbf{t}(\mathbf{s},\mathbf{t})} \left[\frac{\mathbf{x}_j(s',t) - \mathbf{x}(s,t)}{|\mathbf{x}_j(s',t) - \mathbf{x}(s,t)|^3}\right], \tag{S9}$$

with $n_a$ an integer controlling the number of longitudinal cross sections used to define the vector field and $P^\perp_{\mathbf{t}(s,t)} = I - \mathbf{t}\mathbf{t}^\top$ the operator projecting vectors in the plane perpendicular to the unit vector $\mathbf{t}(s,t)$ tangent to the centerline at position $s$ and time $t$. This operator enforces the constraint that the frame is adapted to the centerline, that is $\mathbf{t} \perp \mathbf{M}$. With these constraints, $\mathbf{M}$ can then be used to define the material frame: by defining $\mathbf{d}_1 = \mathbf{M}/|\mathbf{M}|$, $\mathbf{d}_2 = \mathbf{t} \times \mathbf{d}_1$, and using the identification $\mathbf{d}_3 \equiv \mathbf{t}$, the $3 \times 3$ matrix $Q$ whose rows are $\mathbf{d}_1$, $\mathbf{d}_2$ and $\mathbf{d}_3$ is a valid direct orthonormal adapted frame. We then identify the orientation of the directors with the anatomical directions as $\mathbf{d}_1 \equiv \mathbf{d}_{\text{LR}}$, and $\mathbf{d}_2 \equiv -\mathbf{d}_{\text{DV}}$. Example framed curves are shown in Fig. S7d.

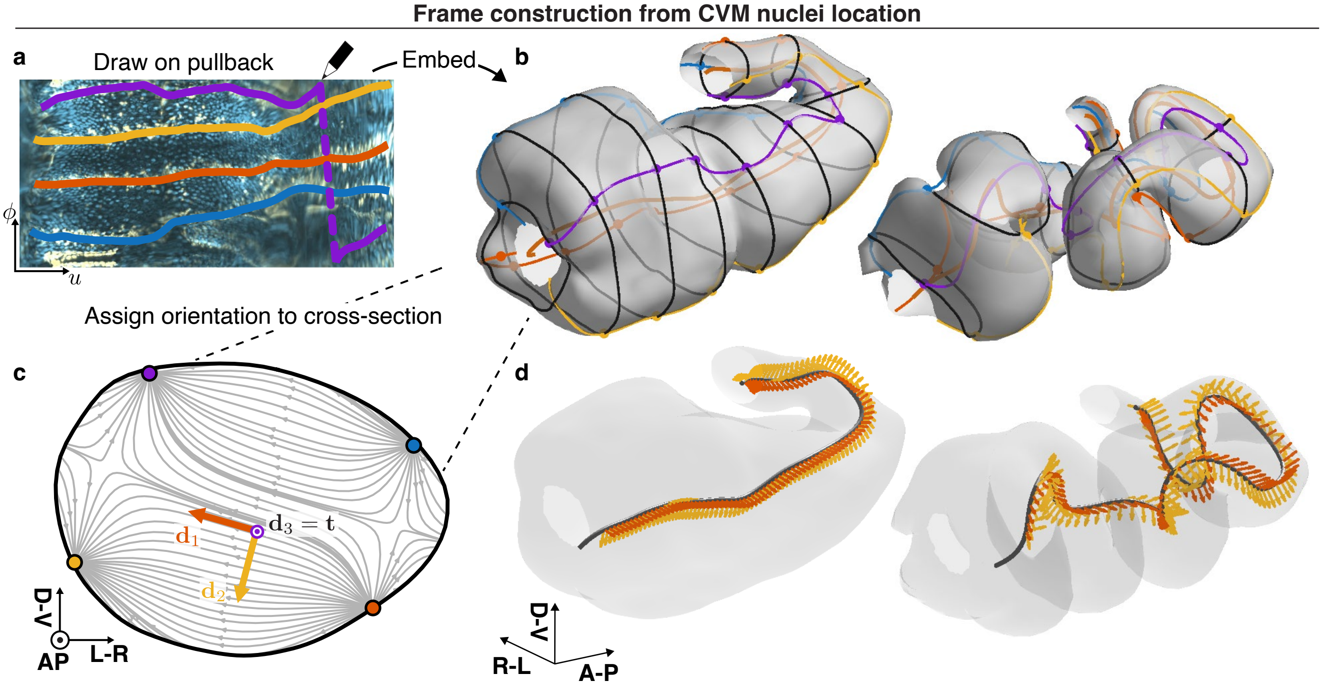


FIG. S7. **Construction of the material frame.** **a.** We identify the locations of the myocyte nuclei by drawing over their projections in the 2D surface texture (blue, orange, yellow and purple curves). **b.** We then embed these positions in 3D, giving us curves in $\mathbb{R}^3$ and enabling us to identify their intersection with coordinate planes (black). **c.** We then use a weighted dipolar field construction to define the cross section orientation (Eq. (S9)). Gray vector field lines represent the **d**. This construction gives us access to a representation of the embryonic midgut as a framed curve, with anatomically-relevant axes.

##### 4. Measures of deformation

The frame vectors are orthonormal and continuously vary along the curve following the kinematic relation [10–12]

$$\mathbf{d}_i' = \boldsymbol{\kappa}(s,t) \times \mathbf{d}_i \tag{S10}$$

where $\boldsymbol{\kappa}$ is the Darboux curvature vector. The components $\kappa^i = \boldsymbol{\kappa} \cdot \mathbf{d}_i$ of $\boldsymbol{\kappa}$ in the material frame, which we collect in $\boldsymbol{\kappa}_L = (\kappa^1, \kappa^2, \kappa^3)$, are the material curvatures, which can be understood as the rotational strains sustained by the rod elements: $\kappa^1$ and $\kappa^2$ induce bending deformations, while $\kappa^3$ corresponds to twisting. The components can be equivalently evaluated by the expression $\kappa^i = \mathbf{d}_i \cdot \mathbf{d}_i'$. The spin vector $\boldsymbol{\omega}(s,t)$ describes the frame's evolution in time

$$\dot{\mathbf{d}}_i = \boldsymbol{\omega}(s,t) \times \mathbf{d}_i. \tag{S11}$$

Similarly as for the curvature, the components $\omega^i = \boldsymbol{\omega} \cdot \mathbf{d}_i$ in the Lagrangian (material-associated) basis made up by the $\mathbf{d}_i$ encode motions of the framed curve, with rotations along $\mathbf{d}_1, \mathbf{d}_2$ corresponding to bending-like degrees of freedom, while motions and deformations along $\mathbf{d}_3$ are associated to twisting. Unlike the curvature, the spin vector is not invariant under rigid-body rotations and is therefore not a measure of deformation. The two are related by the Maurer–Cartan identity, which follows from the equality of mixed derivatives $\partial_s \partial_t \mathbf{d}_i = \partial_t \partial_s \mathbf{d}_i$,

$$\partial_t \boldsymbol{\kappa} = \partial_s \boldsymbol{\omega} - \boldsymbol{\kappa} \times \boldsymbol{\omega}. \tag{S12}$$

which shows that the convected derivative $D_s \boldsymbol{\omega} = \partial_s \boldsymbol{\omega} - \boldsymbol{\kappa} \times \boldsymbol{\omega}$ of the spin vector, rather than $\boldsymbol{\omega}$ itself, measures the rate of deformation.

In the discrete setting, the rod is defined by vertices $\{\mathbf{x}_k\}_{k=1,\ldots,N+1}$ and frames $\{Q_k\}_{k=1,\ldots,N}$, with frames associated to the edges connecting successive vertices. The spin vector can be computed using the matrix logarithm between frames at consecutive timepoints

$$\boldsymbol{\omega}_k = \frac{1}{\Delta t} \log Q_k(t) Q_k(t+\Delta t)^\top, \tag{S13}$$

while the Darboux vector between the frames at successive edges can be found by [13, 14]

$$\boldsymbol{\kappa}_k = \frac{1}{\bar{\ell}_k} \log Q_k Q_{k+1}^\top. \tag{S14}$$

with $\bar{\ell}_k = (\ell_k + \ell_{k+1})/2$ the average of the two successive edge lengths $\ell_k$ and $\ell_{k+1}$, following standard discrete differential geometry practices.

To quantify the relative importance of bending to twisting motions and deformations, we consider different measures. While not a proper deformation measure, a first quantity is the average absolute rotation speed, split into bending and twisting contributions. These read in the continuum setting

$$\bar{\omega}_\mathrm{b}^2 = \frac{1}{L}\int \mathrm{d}s\,(|\omega^1|^2 + |\omega^2|^2), \quad \bar{\omega}_\mathrm{tw}^2 = \frac{1}{L}\int \mathrm{d}s\,|\omega^3|^2. \tag{S15}$$

In the discrete setting, we compute the integrals as $\bar{\omega}_X = (1/L)\sum_k \ell_k |\omega_k^X|$. To measure the overall curvature of the framed curve, we define curvature measures

$$\mathcal{E} = \frac{1}{2}\int \mathrm{d}s\,|\boldsymbol{\kappa}|^2 = \sum_{i=1}^{3} \frac{1}{2}\int \mathrm{d}s\,\kappa_i^2 \tag{S16}$$

which can be split into bending and twisting contributions

$$\mathcal{E}_\mathrm{b} = \frac{1}{2}\int \mathrm{d}s\ \left[(\kappa^1)^2 + (\kappa^2)^2\right] \equiv \mathcal{E}_{\mathrm{b},1} + \mathcal{E}_{\mathrm{b},2}, \tag{S17}$$

$$\mathcal{E}_\mathrm{tw} = \frac{1}{2}\int \mathrm{d}s\,(\kappa^3)^2. \tag{S18}$$

These quantify the geometric complexity of the framed curve rather than its elastic energy, as they do not involve reference curvatures. Discrete integrals are weighted by the Voronoi lengths, $\mathcal{E}_X = (1/2L)\sum_k \bar{\ell}_k (\kappa_k^X)^2$. To assess variation in curvature, we can also consider the time-variation of the curvature components

$$\mu^i = \frac{1}{L}\int \mathrm{d}s\,|\partial_t \kappa^i| \tag{S19}$$

which we discretize similarly as the curvature energies by weighting using $\bar{\ell}_k$. This measure requires taking second derivatives of the data, and is consequently the most sensitive to measurement noise. For all these quantities, to alleviate numerical amplification of experimental noise we compute the integrals after applying a $3 \times 3$ median filter in the (time, arclength) plane which removes isolated frame-to-frame outliers. We plot selected quantities in Fig. S8.

#### V. MEASURING CENTERLINE CHIRALITY AND TOPOLOGY

In this section, we elaborate on the methods used to characterize the geometry and topology of the framed curve. We consider a curve $\gamma \subset \mathbb{R}^3$, framed using the Frenet–Serret basis $(\mathbf{t}, \mathbf{n}, \mathbf{b})$ where it is defined. After presenting a continuous measure of chirality within Frenet–Serret theory, we define a quasi-local measure of chirality more robust to experimental noise which connects to the Writhe, a standard measure of global coiling. We then present some elements of curve topology that are used in the main text to identify the source of chirality generation.

##### 1. Local measure

A key result from Frenet–Serret theory is that a space curve is defined, up to rigid body rotations, by its curvature $\kappa$ and torsion $\tau$, defined by the relationships $\mathbf{t}' = \kappa\mathbf{n}$ and $\mathbf{b}' = -\tau\mathbf{n}$. As above, we use primes to denote differentiation with respect to the curve parameter $s$. While the torsion is a natural measure of chirality, it is discontinuous at inflection points and is very sensitive to noise in nearly-straight portions of the curve.

Here, using the local expansion of the tangent vector [15]

$$\begin{aligned}\mathbf{t}(s+h) &= \mathbf{t}(s) + \mathbf{t}'(s)h + \frac{1}{2}h^2\mathbf{t}''(s) + O(h^3)\\ &= \mathbf{t}(s) + h\kappa\mathbf{n}(s) + \frac{h^2}{2}\left(\kappa'(s)\mathbf{n}(s) - \kappa^2\mathbf{t}(s) + \kappa\tau\mathbf{b}(s)\right) + O(h^3)\end{aligned} \tag{S20}$$

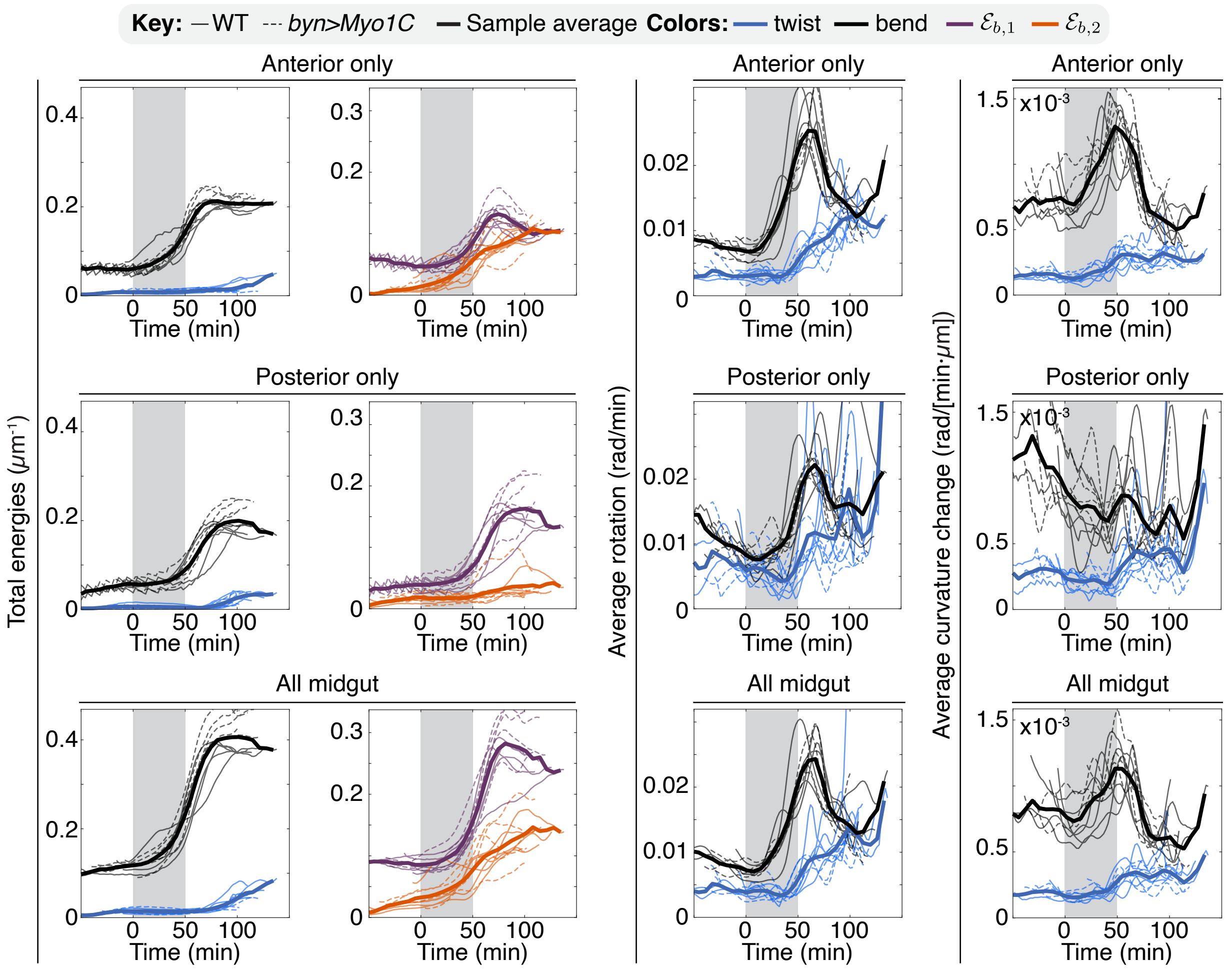


FIG. S8. Decomposition of deformation and motion of the rod. Curvature energies $\mathcal{E}_{\rm b}, \mathcal{E}_{\rm tw}$, bending energies $\mathcal{E}_{\rm b,1}, \mathcal{E}_{\rm b,2}$ integrated over the anterior (top row), posterior (middle) and overall domains (bottom) of the midgut. Average rotation $\bar{\omega}_{\rm b}$, $\bar{\omega}_{\rm tw}$, and variation in curvature $\mu_{\rm bend}$, $\mu_{\rm twist}$ versus time averaged over the anterior (top row), posterior (middle) and overall domains (bottom) of the midgut. Separation between anterior and posterior domains is defined by the posterior-most point of the midgut centerline.

we define a continuous, local measure of chirality by considering the infinitesimal area of the spherical triangle defined by the three points $(\mathbf{t}(s - {\rm d}s), \mathbf{t}(s), \mathbf{t}(s + {\rm d}s))$ [15]. This area will be zero if the curve has no torsion, while in the presence of torsion its sign corresponds to the torsion's. To compute this solid angle, we note that it is given by the area of the spherical triangle on the unit sphere projected in the plane normal to $\mathbf{t}(s)$. This yields a solid angle

$$\begin{aligned} {\rm d}A(s) &= \frac{1}{2}\mathbf{t}(s) \cdot \left( (\mathbf{t}(s + {\rm d}s) - \mathbf{t}(s - {\rm d}s)) \times (\mathbf{t}(s) - \mathbf{t}(s - {\rm d}s)) \right) \\ &= \mathbf{t}(s) \cdot \left[ {\rm d}s\, \mathbf{t}'(s) \times \left( {\rm d}s\, \kappa \mathbf{n}(s) - \frac{{\rm d}s^2}{2} \left( \kappa'(s)\mathbf{n}(s) - \kappa^2 \mathbf{t}(s) + \kappa\tau \mathbf{b}(s) \right) \right) \right] \\ &= \frac{{\rm d}s^3}{2} \kappa^2 \tau. \end{aligned} \tag{S21}$$

As a consistency check, we note that $\kappa$ and $\tau$ both have units of inverse length, such that ${\rm d}A$ is unitless, as solid angles should be. The solid angle then leads to a chirality measure $\chi = 2{\rm d}A/{\rm d}s^3 = \kappa^2\tau$, which has units of inverse length cubed [15].

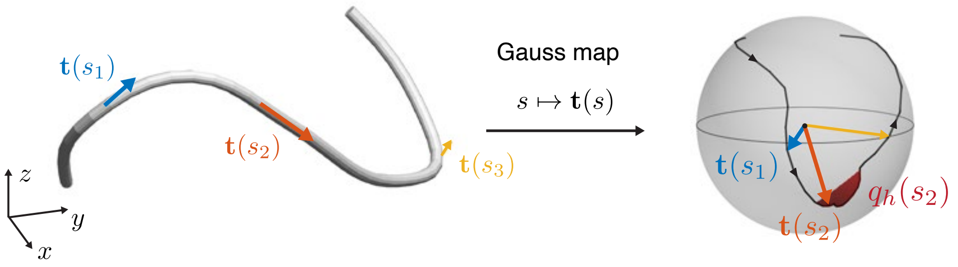


FIG. S9. Quasi-local measure of chirality. We define a quasi-local measure of chirality $q_h(s)$ by considering the signed area swept by the tangent vector to the curve $\mathbf{t}(s)$ on a finite interval $[s-h, s+h]$. If the area is integrated over the whole curve, the resulting measure recovers the Writhe Wr.

##### 2. Quasi-local measure

Evaluating the solid angle above requires taking the second derivative of the tangent vector, or equivalently the third derivative of the curve with respect to its parameter. This measure is thus strongly sensitive to noise, and to mitigate the noise we thus consider a 'weak formulation' of the chiral measure defined above by defining $q_h(s)$ as the solid angle spanned by the curve traced on $S^2$ by $\mathbf{t}(p)$ for $p \in [s-h, s+h]$ and closed by a great circle from $\mathbf{t}(s+h)$ to $\mathbf{t}(s-h)$, illustrated in Fig. S9. The solid angle can be computed by summing the signed area of successive spherical triangles

$$q_h(s) = \int_{s-h}^{s+h} A\left(\mathbf{t}(s-h), \mathbf{t}(p), \mathbf{t}'(p)\,\mathrm{d}p\right). \tag{S22}$$

Numerically, the spherical area is computed using the Van Oosterom–Strackee formula [16]. As $h \to 0$, $q_h(s) \sim h^3\kappa^2\tau/2$ and we recover the local chirality measure. As $h \to L$, the total length of the curve, $q_h(s) \to$ Wr the Writhe of the curve [17]. We use this measure in Main Text Fig. 1 and Fig. S10 to quantify the local chirality of the midguts' centerlines. Numerically, the centerline of length $L$ is represented as a polyline with $N$ equally-spaced nodes, separated by arclength $\Delta s = L/N$, and we sum over the spherical triangle as

$$q_h(s_k) = \sum_{j=k-h/\Delta s}^{k+h/\Delta s} A(\mathbf{t}_{k-h/\Delta s}, \mathbf{t}_j, \mathbf{t}_{j+1}), \tag{S23}$$

with lower and upper limits of the sums adjusted near boundaries. In practice, we use $h = 3\Delta s$ to balance noise suppression with local feature recovery.

##### 3. Link, Writhe, Twist and topological constraint

Consider the framed curve $(\mathbf{x}, Q)$ constructed above, with $\mathbf{d}_1$ the left–right direction and $\mathbf{d}_3 = \mathbf{t}$ the unit tangent. Displacing the centerline by a small distance $\epsilon$ along $\mathbf{d}_1$ defines a second curve $\mathbf{y}(s) = \mathbf{x}(s) + \epsilon\,\mathbf{d}_1(s)$; the two curves $\mathbf{x}$ and $\mathbf{y}$ bound a ribbon, whose geometry and topology are characterized by three quantities.

The *Writhe* Wr depends only on the centerline. It is given by the Gauss double integral

$$\mathrm{Wr} = \frac{1}{4\pi}\int_0^L\int_0^L \mathrm{d}s\,\mathrm{d}s'\,\frac{(\mathbf{t}(s)\times\mathbf{t}(s'))\cdot(\mathbf{x}(s)-\mathbf{x}(s'))}{|\mathbf{x}(s)-\mathbf{x}(s')|^3}, \tag{S24}$$

and can be interpreted as the signed number of self-crossings of the curve, averaged over all viewing directions [17, 18]. It vanishes for planar curves, and its sign indicates the overall handedness of coiling, with $\mathrm{Wr} < 0$ for left-handed coils.

The *Twist* Tw measures the rotation of the frame around the centerline. It is the integral of the tangential component of the Darboux vector, $\kappa^3 = \boldsymbol{\kappa}\cdot\mathbf{t} = (\mathbf{d}_1 \times \mathbf{d}_1')\cdot\mathbf{t}$,

$$\mathrm{Tw} = \frac{1}{2\pi}\int_0^L \mathrm{d}s\,\kappa^3(s), \tag{S25}$$

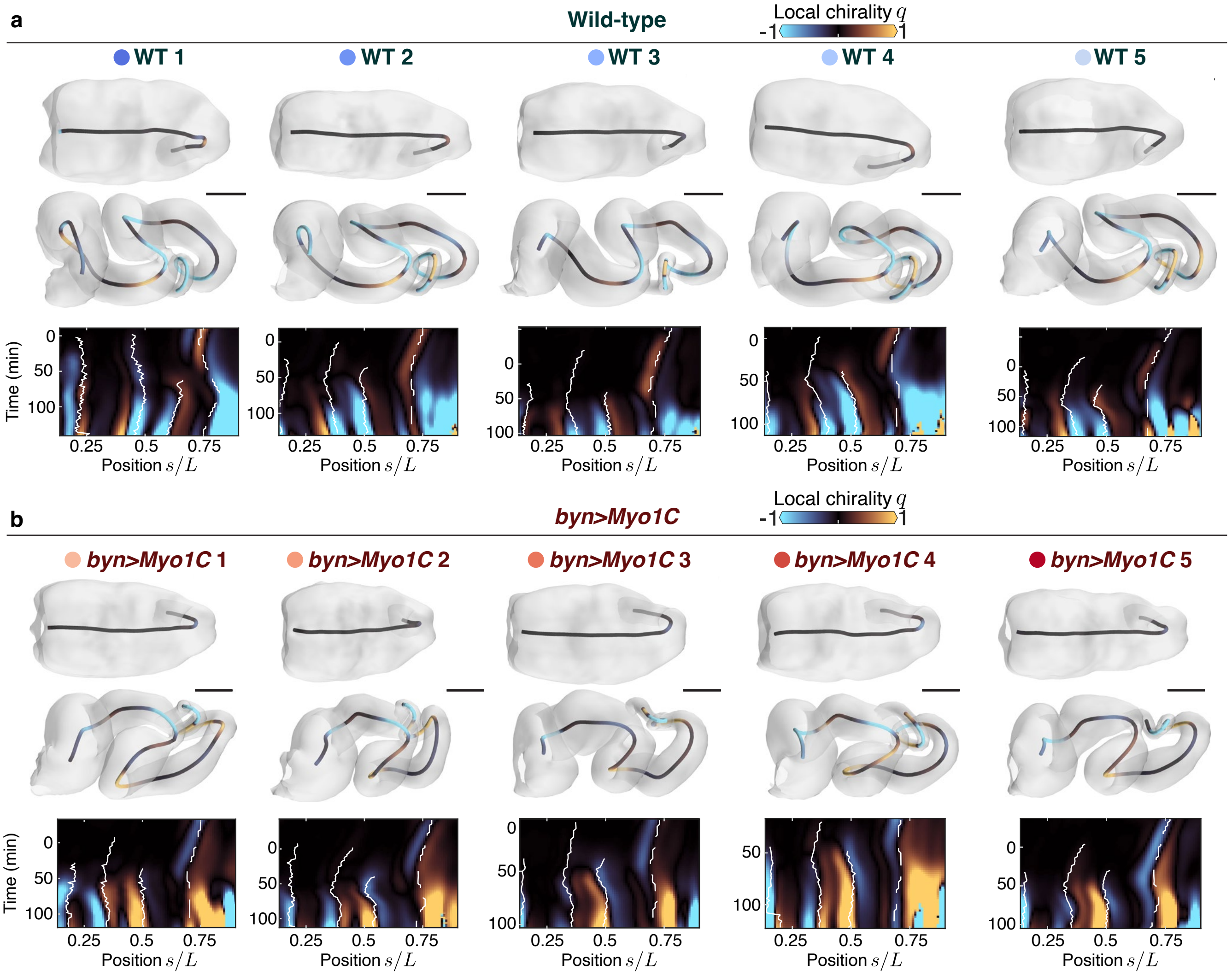


FIG. S10. Local chirality measurement reveal *perverted* coils. We use the chirality metric $q_h$ with $h = 3\Delta s$ to quantify the local chirality of the centerlines for **a.** wild-type and **b.** *byn>Myo1C* samples. 3D renderings shown in ventral view at initial and final times for each sample, scale bars 50 $\mu$m.

that is, of the twist density whose square defines the twisting measure $\mathcal{E}_{\rm tw}$ introduced above [18].

The *Link* Lk is the Gauss linking number of the two curves $\mathbf{x}$ and $\mathbf{y}$, i.e. the number of times they wind around each other. For closed curves, it is an integer and a topological invariant: it can only change if one curve passes through the other. Unlike Wr, both Tw and Lk depend on the frame. Numerically, we compute Twist and Writhe using Algorithms 2a and 1a of Ref. [19].

The Călugăreanu–White–Fuller (CWF) theorem states that for a closed ribbon, the following equality holds between the Link Lk, Writhe Wr and the Twist Tw [17, 18]

$$\mathrm{Lk} = \mathrm{Wr} + \mathrm{Tw} \tag{S26}$$

Writhe and Twist vary continuously as the ribbon deforms, but their sum is fixed as long as the ribbon does not pass through itself: deformations of the ribbon can only trade Writhe for Twist, and vice versa.

The midgut centerline is an open curve. The Gauss integrals defining Wr and Lk remain well defined when evaluated over $[0, L] \times [0, L]$, but for open curves Lk is no longer an integer or a topological invariant, and Eq. (S26) does not hold exactly. We notice, however, that in our data, the sum Wr + Tw of the open ribbon remains approximately constant and close to zero throughout coiling, while the ends of the midgut move and rotate little (Main Text Fig. 1h). This indicates that there exists a closure, fixed in time, which we need not specify, for which the midgut ribbon obeys Eq. (S26) with Lk = 0, the value set by the initially nearly straight and untwisted tube. This effective closure remains valid as long as the midgut does not pass through itself, which self-contact prevents. We thus have

$$\mathrm{Wr} \approx -\mathrm{Tw}. \tag{S27}$$

Equation (S27) is a kinematic identity relating the shape of the centerline to the rotation of the frame. Since the frame is constrained to follow the tissue orientation, the Twist Tw measures an actual twisting deformation of the midgut. With its ends nearly fixed, the midgut can therefore only writhe by twisting its tissue by an equal and opposite amount. Three consequences follow. First, since Lk is conserved, no net twist is injected through the ends: coiling is driven by deformations within the midgut rather than by rotation of its boundaries. Second, the measured Twist need not reflect active twisting of the tissue, since it is the geometric counterpart of the Writhe. Consistent with this idea, twist appears only once the centerline starts to writhe (Main Text Fig. 2e), supporting the absence of active torques in our model. Third, if twisting the tissue is energetically costly, as for any rod that resists torsion [12], the constraint favors configurations that coil locally while limiting the net Writhe, such as perversions [20].

#### VI. VISCOELASTIC COSSERAT ROD DYNAMICS

As the midgut elongates, it becomes increasingly slender and effectively one-dimensional. To provide a biophysical model of the coiling process, we use an overdamped, viscoelastic version of Cosserat rod dynamics [13, 21] as a minimal model of framed curve dynamics. We use the implementation of Cosserat rods in terms of orthogonal matrices from Ref. [13], which we choose over quaternionic representations favored by e.g. Refs. [22, 23]. To approximately account for cellular rearrangement and remodeling processes that happen in the endodermal layer and elsewhere in the tissue [4], we also include Maxwell-like viscoelastic effects with an implementation strategy inspired by Refs. [10, 21].

##### 1. Continuum Cosserat rod theory

Cosserat rod theory provides a framework for the dynamics of framed curves. Throughout this section, we will use the overdot and prime notations to denote derivatives with respect to time and space, namely $\dot{\mathbf{x}} = \partial_t \mathbf{x}$ and $\mathbf{x}' = \partial_s \mathbf{x}$. We first present the construction of the elastic forces from geometry, before detailing the model's dynamics.

###### *1.1. Elastic forces*

Following standard models, the elastic energy of the rod is the sum of a shear/stretch part and a bending/twist part, $E = E_\sigma + E_\kappa$, both written in terms of strains expressed in the material frame. The shear/stretch strain $\boldsymbol{\sigma}_L = Q\mathbf{x}' - \mathbf{e}^3$, with $\mathbf{e}^3 = (0, 0, 1)^\top$ and the derivative $\mathbf{x}'$ taken with respect to the *rest* arclength, vanishes for an unstretched rod whose frame is *adapted*, $\mathbf{d}_3 = \mathbf{t}$. The bending/twist strain is the deviation of the material curvatures $\boldsymbol{\kappa}_L$ from their reference values $\boldsymbol{\kappa}_L^0$. The two energies read

$$E_\sigma = \frac{1}{2} \int \mathrm{d}s \, \boldsymbol{\sigma}_L \cdot \mathbf{S} \, \boldsymbol{\sigma}_L, \qquad E_\kappa = \frac{1}{2} \int \mathrm{d}s \, \left(\boldsymbol{\kappa}_L - \boldsymbol{\kappa}_L^0\right) \cdot \mathbf{B} \left(\boldsymbol{\kappa}_L - \boldsymbol{\kappa}_L^0\right), \tag{S28}$$

and yield the constitutive relations for the internal force $\mathbf{n}$ and internal couple $\mathbf{m}$, expressed in the material frame,

$$\mathbf{n}_L = \mathbf{S} \, \boldsymbol{\sigma}_L, \qquad \mathbf{m}_L = \mathbf{B} \left(\boldsymbol{\kappa}_L - \boldsymbol{\kappa}_L^0\right). \tag{S29}$$

Compared to Kirchhoff rods, Cosserat rods have an additional shearing degree of freedom, which materializes as the difference between $\mathbf{d}_3$ and the centerline tangent $\mathbf{t}$; a large shear stiffness acts as a soft constraint keeping the frame adapted. The shear/stretch stiffness matrix $\mathbf{S}$ and bending/twist stiffness matrix $\mathbf{B}$ are parametrized as diagonal matrices

$$\mathbf{S} = \begin{pmatrix} S_1 & & \\ & S_2 & \\ & & S_3 \end{pmatrix}, \qquad \mathbf{B} = \begin{pmatrix} B_1 & & \\ & B_2 & \\ & & B_3 \end{pmatrix}. \tag{S30}$$

In our simulations, we simplify the geometry by assuming $S_1 = S_2$ and $B_1 = B_2$, take $S_1$ and $S_3$ to be sufficiently large to assume an unshearable rod of prescribed (time-dependent) length, and vary the ratio $B_3/B_1$. Even using this rough approximation, we find that the visco-elastic dynamics below are able to recover experimental shape dynamics.

##### 1.2. Overdamped viscoelastic dynamics

The midgut is immersed in a complex viscous medium and contacts itself at later stages. To obtain an effective dynamical model of the midgut deformation, we assume simple overdamped dynamics for both displacements and rotational degrees of freedom. More specifically, we assume that the deformations of the rod induce internal forces $\mathbf{n}$ and couples $\mathbf{m}$ deriving from the elastic energy above. We additionally allow external forces $\mathbf{f}^{\text{ext}}$ and couples $\mathbf{m}^{\text{ext}}$ acting on the rod. Overdamped momentum balance in the Eulerian frame reads

$$\gamma_L \dot{\mathbf{x}} = \partial_s \mathbf{n} + \mathbf{f}^{\text{ext}}, \tag{S31}$$

$$\gamma_R \boldsymbol{\omega} = \partial_s \mathbf{m} + \mathbf{x}' \times \mathbf{n} + \mathbf{m}^{\text{ext}}. \tag{S32}$$

with $\boldsymbol{\omega}$ the spin vector of the frame, which updates the material frame via Eq. (S11). We explicitly write down $\mathbf{n}$ and $\mathbf{m}$ in the discrete setting in the next section.

Given the observation that dissected midguts partially keep their shape (data not shown) and the presence of internal rearrangements of the endoderm [4], we assume that there exist viscoelastic effects that update the reference condition. We thus implement a Maxwell-like viscoelastic update rule for the reference curvature $\boldsymbol{\kappa}^0$: the rest state relaxes towards the instantaneous curvature at a rate $\eta$, so that sustained bending and twisting are progressively 'written in' as new rest shape while transient deformations remain elastic. Concretely, the reference curvature obeys [24]

$$\partial_t \boldsymbol{\kappa}^0 = \eta\,(\boldsymbol{\kappa} - \boldsymbol{\kappa}^0), \tag{S33}$$

which is integrated using an implicit Euler scheme in the discrete dynamics (Eq. (S43)). The single parameter $\eta$ interpolates between the purely elastic limit ($\eta \to 0$, fixed rest shape) and a rapidly-yielding limit ($\eta \to \infty$, where the rod's reference configuration is vanishingly close to the current configuration). The stretching and shearing responses are kept purely elastic.

#### 2. Discretization of the Cosserat rod

Following Ref. [13], we now present the discretized version of the Cosserat rod framework that we will use to simulate midgut coiling. The centerline of the rod is represented numerically as a chain of $N+1$ nodes $\mathbf{x}_k \in \mathbb{R}^3$, $k = 1, \ldots, N+1$, connected by $N$ edges that carry the material frames $Q_k$, whose rows are the directors as above. We denote by $\ell_k = |\mathbf{x}_{k+1} - \mathbf{x}_k|$ the length of the $k$-th edge, by $\ell_k^0$ its rest length, and by $\mathbf{t}_k = (\mathbf{x}_{k+1} - \mathbf{x}_k)/\ell_k$ its unit tangent; for high extensional stiffness, $\ell_k \approx \ell_k^0$. In continuous time, Eq. (S11) reads

$$\partial_t Q = -Q\,\text{skew}(\boldsymbol{\omega}). \tag{S34}$$

We adapt the framework of Ref. [13] to the overdamped case to define internal forces and couples and discretize the time dynamics; note that our definition of the material frame differs slightly from theirs, so that some relations below differ from the reference. The key idea is that in the material frame of the rod, forces and torques are readily written in terms of curvature and frame vectors, which can then be expressed in the lab (simulation) frame by a simple change of variable to generate physically-consistent dynamics. Explicitly, to switch between a vector $\mathbf{v}_L$ expressed in the material (Lagrangian) frame and $\mathbf{v}$ in the lab frame, we use the relations

$$\mathbf{v} = Q^\top \mathbf{v}_L \iff \mathbf{v}_L = Q\mathbf{v} \tag{S35}$$

The discrete Darboux vectors $\boldsymbol{\kappa}_k$ are computed from successive frames using Eq. (S14), with $\bar{\ell}_k$ the length of the Voronoi domain of the $(k+1)$-th vertex. Introducing $\varepsilon_k = \ell_k/\ell_k^0$ the extensional elongation of the $k$-th edge, and $e_k = \bar{\ell}_k/\bar{\ell}_k^0$ the dilatation of its Voronoi domain, with $\bar{\ell}_k^0$ the rest Voronoi length, the internal shear/stretch force $\mathbf{n}_k$ and bending/twisting couple $\mathbf{m}_k$ are expressed in the lab frame as

$$\mathbf{n}_k = Q_k^\top\,\mathbf{S}\,(\varepsilon_k\,Q_k\mathbf{t}_k - \mathbf{e}^3), \qquad \mathbf{m}_k = \frac{1}{e_k^3}\,Q_k^\top\,\mathbf{B}\,Q_k\,(\boldsymbol{\kappa}_k - \boldsymbol{\kappa}_k^0), \tag{S36}$$

where the $e_k^{-3}$ factor rescales the bending response by the local Voronoi dilatation [13], and $\mathbf{e}^3$ is defined as above. The constitutive matrices $\mathbf{S}$ and $\mathbf{B}$ are diagonal only in the material frame, so each stress is formed by rotating the relevant strain into the material frame (left-multiplication by $Q_k$), applying the diagonal stiffness, and rotating the resulting force or couple back to the lab frame ($Q_k^\top$). The nodal forces $\mathbf{f}_k$ and edge torques $\boldsymbol{\Gamma}_k$ in the lab frame are then assembled from these stresses as

$$\mathbf{f}_k = \Delta\,[\mathbf{n}_k] \tag{S37}$$

$$\boldsymbol{\Gamma}_k = \Delta\,[\mathbf{m}_k] + \mathcal{A}\,[\bar{\ell}_k^0\,\boldsymbol{\kappa}_k \times \mathbf{m}_k] + \ell_k^0\,\mathbf{n}_k \times \mathbf{t}_k \tag{S38}$$

where the discrete difference operator $\Delta[\cdot] : \mathbb{R}^{3\times N} \to \mathbb{R}^{3\times(N+1)}$ and discrete averaging operator $\mathcal{A}[\cdot] : \mathbb{R}^{3\times N} \to \mathbb{R}^{3\times(N+1)}$ acting on $N$-uples of vectors in $\mathbb{R}^3$ are defined by

$$\Delta[\mathbf{x}_{k=1,\ldots,N}] = \begin{cases} \mathbf{x}_1 & \text{if } j=1 \\ \mathbf{x}_j - \mathbf{x}_{j-1} & \text{if } 1<j\leq N \\ -\mathbf{x}_N & \text{if } j=N+1 \end{cases}, \qquad \mathcal{A}[\mathbf{x}_{k=1,\ldots,N}] = \begin{cases} \mathbf{x}_1/2 & \text{if } j=1 \\ (\mathbf{x}_j + \mathbf{x}_{j-1})/2 & \text{if } 1<j\leq N \\ \mathbf{x}_N/2 & \text{if } j=N+1 \end{cases}. \tag{S39}$$

The forces and torques lead to the overdamped time derivatives of the position and frame

$$\frac{\mathrm{d}\mathbf{x}_k}{\mathrm{d}t} = \frac{1}{\gamma_L}(\mathbf{f}_k + \mathbf{f}_k^{\mathrm{ext}}) \tag{S40}$$

$$\frac{\mathrm{d}Q_k}{\mathrm{d}t} = Q_k \,\mathrm{skew}\left[(\mathbf{\Gamma}_k + \mathbf{m}_k^{\mathrm{ext}})/\gamma_R\right] \tag{S41}$$

where $\gamma_L$ and $\gamma_R$ are the translational and rotational drag coefficients. To integrate these equations we use a standard four-stage Runge-Kutta (RK4) scheme applied jointly to the vertex positions and the frames. Within each stage the vertex positions advance by an explicit Euler substep, while the frame increment $\boldsymbol{\omega}\,\Delta t$ is applied through the matrix exponential computed using Rodrigues' formula,

$$Q_k(t+\Delta t) = Q_k(t)\,\exp\left(\Delta t\,\mathrm{skew}(\boldsymbol{\omega}_k)\right), \tag{S42}$$

so that each $Q_k$ remains a rotation matrix at every stage; the rows are re-normalized after each update to suppress numerical drift. The four stage increments are then combined with the usual RK4 weights $(1,2,2,1)/6$.

Finally, the viscoelastic rest curvatures are updated with an implicit (backward-Euler) step, which discretizes Eq. (S33) and is unconditionally stable in $\eta\Delta t$,

$$\boldsymbol{\kappa}_k^0(t+\Delta t) = \frac{\boldsymbol{\kappa}_k^0(t) + \eta\Delta t\,\boldsymbol{\kappa}_k(t+\Delta t)}{1+\eta\Delta t}, \tag{S43}$$

with the current curvature $\boldsymbol{\kappa}_k(t+\Delta t)$ estimated from the updated frames using Eq. (S14).

##### 3. Growth-rate field from data

The rod elongates according to a prescribed growth field $g(\bar{s},t)$, with $\bar{s}$ the fractional arclength defined below, that sets the rate of change of the rest lengths, such that $\dot{\ell}^0 = g$. Rather than imposing an idealized growth law, we measure this field directly from the reconstructed organ so that the simulated rod reproduces the observed, spatially non-uniform elongation of the midgut.

###### *3.1. Construction from data*

At each experimentally measured timepoint $t_a$ ($a = 0,\ldots,K$), the `TubULAR` reconstruction (Sec. III) provides the centerline as a polyline with vertices $\mathbf{x}_k(t_a)$. We form the edge lengths, cumulative arclengths, and total length

$$\ell_k(t_a) = |\mathbf{x}_{k+1}(t_a) - \mathbf{x}_k(t_a)|, \qquad s_k(t_a) = \sum_{k'<k} \ell_{k'}(t_a), \qquad L(t_a) = \sum_k \ell_k(t_a), \tag{S44}$$

and define the fractional arclength coordinate $\bar{s}_k(t_a) = s_k(t_a)/L(t_a) \in [0,1]$. Using $\bar{s}$ rather than the physical arclength provides a common material-like coordinate along which growth can be compared across timepoints despite the overall elongation.

Because the vertex positions are noisy, we first smooth the edge-length profile in space with a few iterations ($n_{\mathrm{sm}} = 4$) of discrete diffusion,

$$\ell_k \leftarrow \ell_k + \tfrac{1}{2}\left(\ell_{k-1} + \ell_{k+1} - 2\ell_k\right), \tag{S45}$$

which suppresses high-wavenumber noise, at the mild cost of slightly reducing the overall magnitude. We then form the per-frame length increments $\Delta_a\ell_k = \ell_k(t_{a+1}) - \ell_k(t_a)$ and represent them as a function of the fractional arclength by linear interpolation $\Delta_a\ell(\bar{s})$, with flat extrapolation at the ends.

Finally, we map the $K$ measured intervals onto the simulation time axis $[0,T]$, assigning each interval a duration $\Delta t_{\rm data} = T/K$, and rescale lengths by $L$ to obtain the non-dimensional growth-rate field

$$g(\bar{s},t) = \frac{\Delta_a \ell(\bar{s})}{L\,\Delta t_{\rm data}}, \qquad t \in [t_a, t_{a+1}], \tag{S46}$$

which we store as a bilinear interpolant on the grid $(t,\bar{s}) \in [0,T] \times [0,1]$ (with flat extrapolation) (Fig. S11a). By construction, integrating $g$ over the simulation reproduces the measured cumulative elongation, $\int_0^T g(\bar{s},t)\,\mathrm{d}t = \Delta\ell_{\rm tot}(\bar{s})/L$.

##### *3.2. Application in the simulation*

At each timestep we recompute the current fractional arclength $\bar{s}_k$ of every edge (which changes as the rod grows and is remeshed), sample the field at the current simulation time, and update the rest lengths additively,

$$\ell_k^0(t+\Delta t) = \ell_k^0(t) + g(\bar{s}_k,t)\,\Delta t. \tag{S47}$$

Because growth is prescribed in the fractional (material) coordinate, each material segment samples the same portion of the growth program as the rod elongates and is refined by remeshing. The interpolated circumradius field $r(s,t)$ used for self-contact (Sec. VI 6) uses the same procedure, interpolating the measured cross-sectional radius as a function of $(\bar{s},t)$.

#### 4. Boundary conditions

We use the experimentally observed endpoint positions and orientations to construct time-dependent boundary conditions that interpolate between observed time points. In more detail, the locations of the first and last simulation nodes $\mathbf{x}_1(t)$ and $\mathbf{x}_{N+1}(t)$ are constrained to follow the experimental trajectories $\mathbf{x}(\bar{s}=0,t)$, $\mathbf{x}(\bar{s}=1,t)$, linearly interpolating between measured time points. At each simulation time point, their positions are prescribed and influence the forces felt by the rest of the system. The orientations of the first and last frames $Q_1(t)$ and $Q_N(t)$ are set in a similar way by the measured frames $Q(\bar{s},t)$ at $\bar{s}=0,1$. Note that the experimental data is cropped a little at the anterior and posterior ends to avoid numerical artifacts from frame constructions and geometric inaccuracies from the artificial endcap closure procedure.

#### 5. Remeshing

To maintain a fine-enough discretization throughout the inhomogeneous growth process, we implement an adaptive remeshing scheme, in which rod segments are split when they get too long. More specifically, every 5 timesteps we test each edge against a threshold length $\ell_c$ (taken as twice the average initial edge length), and insert a new node at the midpoint of every edge that exceeds it. When an edge of rest length $\ell_k^0$ is split, each of the two children is assigned half the parent rest length, $\ell_k^0/2$, and both inherit the parent's material frame $Q_k$ and its elastic moduli.

The rest curvatures are assigned so as not to inject spurious strain: the first (anterior) child retains the parent edge's reference curvature $\boldsymbol{\kappa}_k^0$, while the newly inserted edge is initialized straight, with vanishing reference curvature $\boldsymbol{\kappa}^0 = \mathbf{0}$ (Fig. S11c). Because the two children share the same frame, the newly created interior node carries zero discrete curvature at insertion, so remeshing leaves the elastic energy unchanged and the subsequent viscoelastic relaxation (Eq. (S43)) sets the new rest curvature over time.

#### 6. Self-contact

We define the circumradius $r(s,t)$ of a point $\mathbf{x}(s,t)$ on the centerline as the distance from $\mathbf{x}(s,t)$ to the closest point on the surface of the gut

$$r(s,t) = \min_{u,\phi} |\mathbf{X}(u,\phi,t) - \mathbf{x}(s,t)|. \tag{S48}$$

We use the circumradius as the effective tube radius carried by each node, so that two nodes repel once their tubes overlap (Fig. S11b). To avoid spuriously flagging nodes that are simply consecutive along the (thick) rod, we

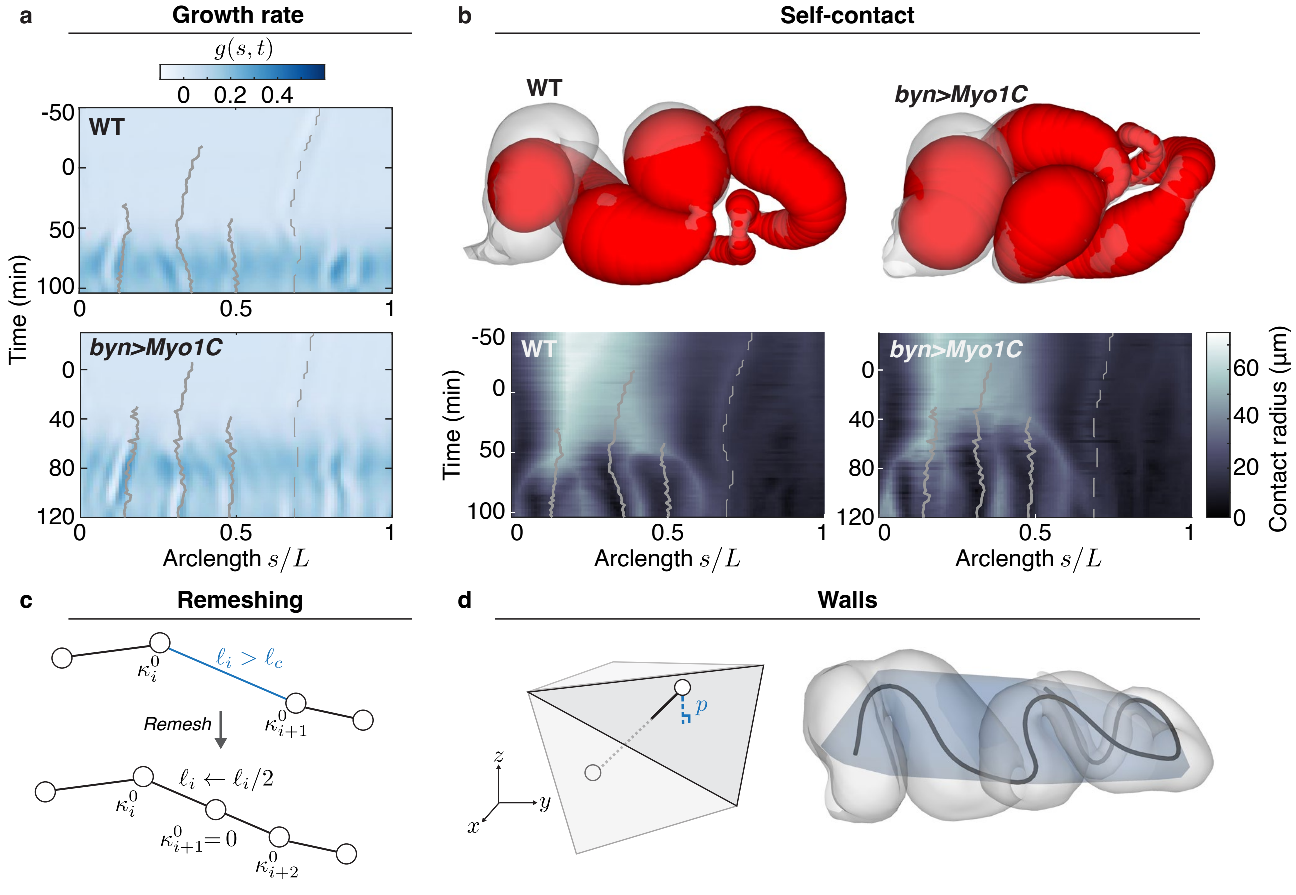


FIG. S11. Experimentally-constrained rod dynamics. **a.** Measured growth field $g(\bar{s}, t)$ for representative WT and *byn>Myo1C* samples. Note that the centerline growth rate is different than the surface elongation. **b.** Self-contact is implemented using the circumradius to define spheres of exclusion for the thick rod that models the midgut. **c.** Illustration of the remeshing procedure, leading to a new segment with no preferred curvature at the new node. **d.** Anatomical confinement is implemented as linear springs confining the simulation nodes inside a polyhedron derived from experimental node positions.

distinguish along-rod proximity from self-contact by their arclength separation: a pair $(i, j)$ is added to the contact neighbor list only if their arclength separation exceeds $r_i + r_j$ and their Euclidean distance is below a cutoff $\propto (r_i + r_j)$. Crucially, this allows deeply interpenetrating segments to not drop out of the other's list and tunnel through each other. This list is rebuilt every 5 timesteps alongside the wall neighbor lists.

For each node $i$ we then take its closest admitted neighbor $j$ and define the overlap $d = \max(r_i + r_j - |\mathbf{x}_i - \mathbf{x}_j|, 0)$. The pair repels along the line of centers according to

$$\mathbf{f}_{i \to j} = -K_1 \, d \, (1 + K_2 \, d) \, \hat{\mathbf{r}}_{ij}, \tag{S49}$$

with $\hat{\mathbf{r}}_{ij}$ the unit vector from $\mathbf{x}_i$ to $\mathbf{x}_j$ and $\mathbf{f}_{j \to i} = -\mathbf{f}_{i \to j}$. In practice we use a purely linear repulsion (contact stiffness $K_1 = 3 \times 10^2$ in non-dimensional units, $K_2 = 0$); the two clamped endpoints are excluded from self-contact forces. Simulations appear not sensitive to the precise form of the interaction parameters, provided the repulsion force is able to overcome internal mechanical forces.

##### 7. Anatomical confinement

We use the convex hull of the downsampled centerline data to define a convex polyhedron defining the anatomical cavity. The convex hull is initially defined in terms of a triangulation with the $i$-th face defined by the triple of points $(V_1^i, V_2^i, V_3^i)$. We convert this triangulation to a half-space representation: each halfspace is defined by the inequality

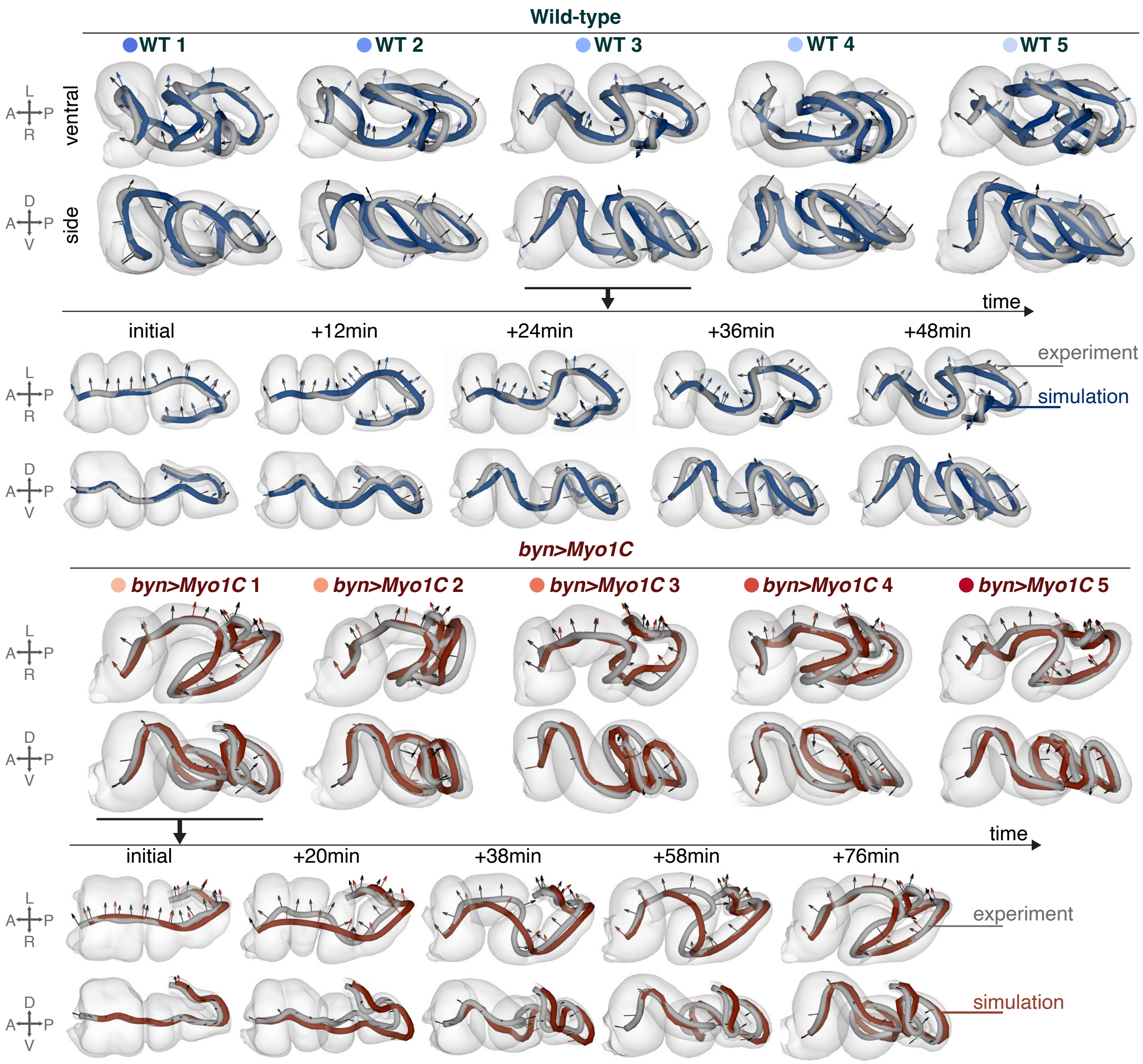


FIG. S12. Simulations of midguts using experimental constraints recovers the dynamics and morphology of individual samples. Samples WT 3 and *byn>Myo1C* 1 illustrate the recovery of coiling dynamics. $B_3/B_{1,2} = 1$, $\eta T = 10$, $T/\tau_E = 3$.

$\boldsymbol{\nu}_i \cdot \mathbf{x} \leq d_i$, with $\boldsymbol{\nu}_i$ the outward-pointing normal and $d_i \in \mathbb{R}$ a real parameter. The orientation is defined by using the position of the centroid of the bounding polyhedron $\mathbf{x}_c$, such that $\boldsymbol{\nu}_i \cdot \mathbf{x}_c \leq d_i$.

The force from the boundary felt at position $\mathbf{x}$ is only non-zero if $\mathbf{x}$ is outside the boundary and given by

$$\mathbf{f}_{\text{wall}}(\mathbf{x}) = -W_1 p(1 + W_2 p)\boldsymbol{\nu} \tag{S50}$$

with $p = \boldsymbol{\nu} \cdot \mathbf{x} - d \geq 0$ the distance to the closest face, defined by the pair $(\boldsymbol{\nu}, d)$. In practice we use a stiff linear restoring force ($W_1 = 10^4$ in non-dimensional units, $W_2 = 0$), which is sufficient to confine the rod to the cavity; the two clamped endpoints are exempted from the boundary force. The quadratic stiffening term $W_2$ is retained in the implementation as an option but is not needed for the simulations reported here (Fig. S11d).

For computational efficiency, as the confining polyhedra feature around a hundred faces, we maintain for each discretization point of the rod a list of nearby faces (those within a cutoff distance $d_c$, taken as $d_c = 0.1\, L_0$), which is rebuilt every few (in practice 5) time steps together with remeshing and self-contact bookkeeping.

##### 8. Parameter sweeps

We non-dimensionalize the governing equations by rescaling lengths by the initial length $L_0$ of the experimental centerline (Eq. (S51)), so that the rescaled rod has unit length, and time by the characteristic bending relaxation timescale $\tau_E = \gamma_R L_0^2/B_1$. In these units we set $L_0 = 1$, $\gamma_L = \gamma_R = 1$, and $B_1 = 1$, so that a total simulation time $T = 1$ corresponds to one bending relaxation time.

Ignoring self-contact and assuming stiff walls, the physics of our growing viscoelastic rods is then controlled by a small set of dimensionless numbers: the twist-to-bend ratio $B_3/B_1$; the dimensionless viscoelastic rate ('Fluidity', or inverse Deborah number) $\eta\tau_E$, which sets how quickly deformations are written into the rest shape; the simulation time $T/\tau_E$, which sets how fast elastic effects affect the rod; and the dimensionless growth, encoded by the prescribed elongation-rate field $g$ extracted from the surface data. The remaining stiffnesses are set large enough to enforce constraints: the stretch and shear rigidities $S_{1,2}L_0^2/B_1 = S_3 L_0^2/B_1 = 10^3$ are large, placing the rod in the near-inextensible, unshearable limit, and the wall and contact stiffnesses are likewise taken large enough to enforce confinement and non-penetration. We sweep $T/\tau_E$, $B_3/B_1$ and $\eta\tau_E$, each over roughly 0.1–30, 0.3–10 and 0.1–$10^3$ respectively. See Table S 2 for all parameter values.

To determine optimal values of the swept parameters, we assess the quality of the simulation by defining measures of both position and orientation reconstruction.

###### *8.1. Resampling onto the experimental grid*

The reference (experimental) configuration is the discrete framed curve $\big(\mathbf{x}_k^{\mathrm{ref}}(t_a), Q_k^{\mathrm{ref}}(t_a)\big)$, sampled at the $N_t$ experimental time points $t_a$, $a = 1, \ldots, N_t$, and at $N_{\mathrm{ref}}$ nodes along the centerline, $k = 1, \ldots, N_{\mathrm{ref}}$, retaining every second node of the reconstructed centerline. Its initial length

$$L_0 = \sum_{k=1}^{N_{\mathrm{ref}}-1} \left|\mathbf{x}_{k+1}^{\mathrm{ref}}(t_1) - \mathbf{x}_k^{\mathrm{ref}}(t_1)\right| \tag{S51}$$

sets the unit of length used throughout the simulations.

The simulation output, saved at $N_{\mathrm{sim}}$ time points on $[0, T]$, is linearly resampled onto the experimental grid in two stages: first in the normalized time $\bar{t} = t/T \in [0, 1]$, then in the fractional arclength $\bar{s}$ (Sec. VI 3), computed from each configuration's own cumulative arclength. Frames, which are defined on edges, are placed at edge midpoints. We denote the resampled fields by $\tilde{\mathbf{x}}_k^{\mathrm{sim}}(t_a)$ and $\tilde{Q}_k^{\mathrm{sim}}(t_a)$. Since the total length of the rod is prescribed by the stiff elongation constraint, comparing configurations at equal $\bar{s}$ isolates differences in shape and orientation.

###### *8.2. Position error*

We measure the position error as the root-mean-square error averaged over time points,

$$E_{\mathrm{pos}} = \frac{1}{N_t\sqrt{N_{\mathrm{ref}}}} \sum_{a=1}^{N_t} \left[\sum_{k=1}^{N_{\mathrm{ref}}} \left|\mathbf{x}_k^{\mathrm{ref}}(t_a) - \tilde{\mathbf{x}}_k^{\mathrm{sim}}(t_a)\right|^2\right]^{1/2}, \tag{S52}$$

in units of $L_0$. In these units, an error of about 6% corresponds to one mean midgut radius, and 11% to one maximal radius.

###### *8.3. Orientation error*

To compare orientations, we measure the rotation of the frame about the body axis. Let $\hat{\mathbf{e}}$ be the unit vector along the body axis (the lab $x$ axis), and $(\hat{\mathbf{e}}_1, \hat{\mathbf{e}}_2)$ an orthonormal basis of the plane normal to it. For a frame vector $\mathbf{d}$, we define its azimuth $\psi$ and the norm $\rho$ of its projection in this plane,

$$\psi(\mathbf{d}) = \operatorname{atan2}(\mathbf{d}\cdot\hat{\mathbf{e}}_2,\, \mathbf{d}\cdot\hat{\mathbf{e}}_1), \qquad \rho(\mathbf{d}) = \left[(\mathbf{d}\cdot\hat{\mathbf{e}}_1)^2 + (\mathbf{d}\cdot\hat{\mathbf{e}}_2)^2\right]^{1/2}. \tag{S53}$$

The azimuthal error of the left–right vector $\mathbf{d}_1$ is

$$\Delta\psi = \left|\left[\psi(\mathbf{d}_1^{\mathrm{sim}}) - \psi(\mathbf{d}_1^{\mathrm{ref}})\right]_{(-\pi,\pi]}\right|, \tag{S54}$$

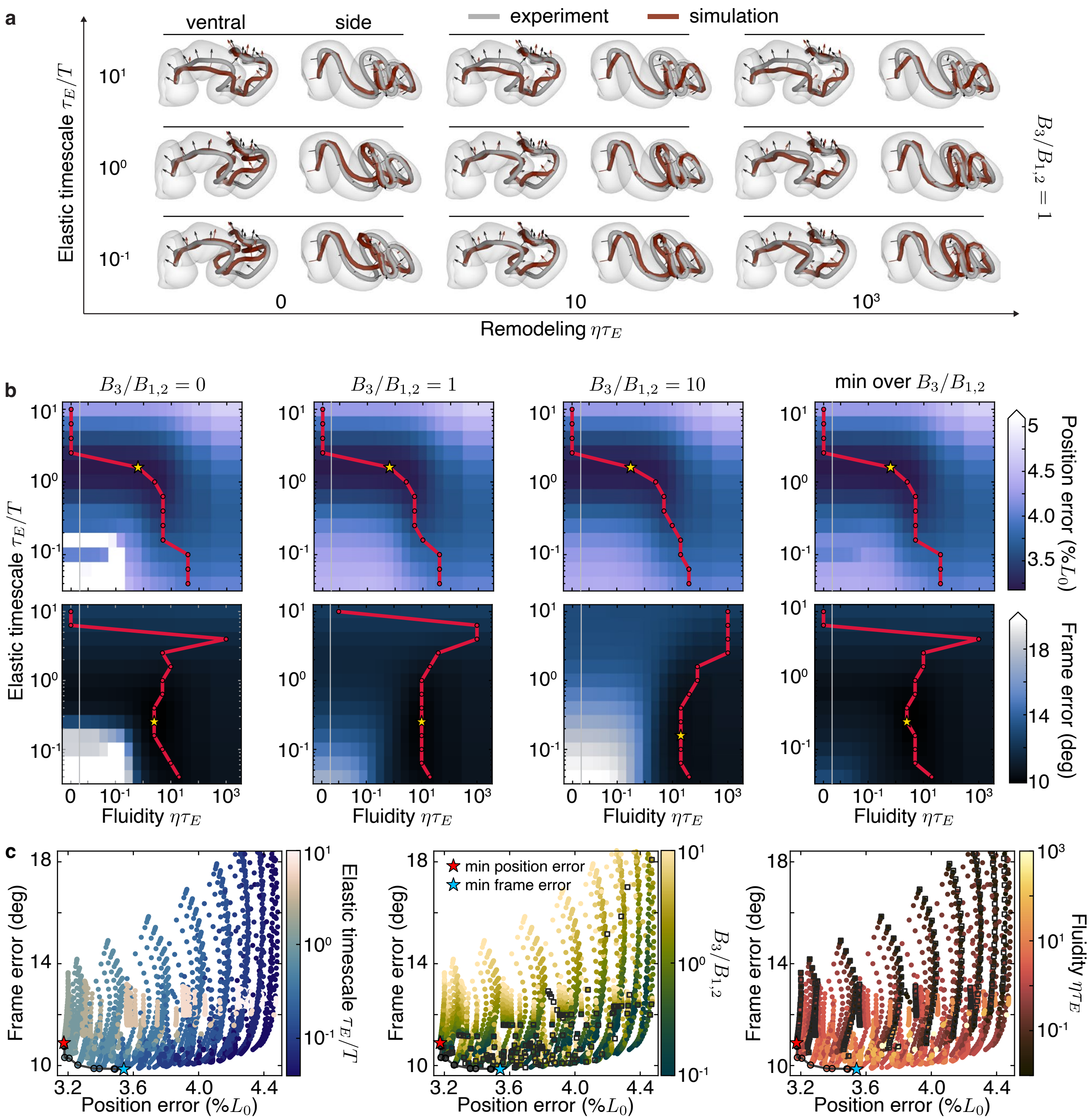


FIG. S13. Parameter sweep shows robust recovery of experimental dynamics. **a**. Selected simulation results, shown at final timepoints and compared to experimental centerline and frame. Choice of parameter weakly affects the final global morphology. **b**. Position and Frame errors versus Fluidity $\eta\tau_E$ and Elastic timescale $\tau_E/T$ for selected values of $B_3/B_1$ and minima over $B_3/B_1$. Red line indicates the argmin over Fluidity. Stars indicate minimum. **c**. Pareto plots showing optimal parameter ranges for joint minimization of Frame and Position errors. Line indicates Pareto-optimal points, blue star minimum of Frame error, red star minimum of Position error.

where $[\cdot]_{(-\pi,\pi]}$ wraps angles into $(-\pi, \pi]$. This measure is ill-conditioned where $\mathbf{d}_1$ aligns with $\hat{\mathbf{e}}$, which happens when $\rho \to 0$. With $\rho_{\min} = \min\left(\rho(\mathbf{d}_1^{\text{sim}}), \rho(\mathbf{d}_1^{\text{ref}})\right)$, we therefore report the conditioning-weighted root-mean-square error as the frame error

$$E_{\text{frame}} = \left( \frac{\sum_{a,k} \rho_{\min}^2 \, \Delta\psi^2}{\sum_{a,k} \rho_{\min}^2} \right)^{1/2}, \tag{S55}$$

| Parameter | Simulation Value |
|---|---|
| Simulation time $T$ (swept) | $T/\tau_E \in [0.1, 30]$ (swept) |
| Bending stiffness $B_{1,2}$ | 1 |
| Twisting stiffness $B_3$ | $B_3/B_1 \in [0.3, 10]$ (swept) |
| Viscoelastic rate $\eta$ | $\eta\tau_E \in [0.1, 10^3]$ (swept) |
| Shear stiffness $S_{1,2}$ | $10^3$ |
| Stretch stiffness $S_3$ | $10^3$ |
| Translational drag $\gamma_L$ | 1 |
| Rotational drag $\gamma_R$ | 1 |
| Contact stiffness $K_1$ | $3 \times 10^2$ |
| Contact stiffening $K_2$ | 0 |
| Wall stiffness $W_1$ | $10^4 > S_3$ |
| Wall stiffening $W_2$ | 0 |
| Remesh threshold $\ell_c$ | $2\,\ell_{\text{init}}$ |
| Wall/contact cutoff $d_c$ | 0.1 |
| Timestep $\Delta t$ | $3 \times 10^{-4}\,\ell_{\min}$ |

TABLE S 2. Parameters for viscoelastic Cosserat rod simulations, in non-dimensional units ($L_0 = 1$, $\gamma_L = \gamma_R = 1$, $B_1 = 1$). Here $\ell_{\min}$ is the smallest initial edge length and $\ell_{\text{init}}$ the initial average edge length. Values quoted as "swept" are varied across the parameter study.

which down-weights ill-conditioned entries.

##### 9. Effect of confinement and boundary orientation

To understand the effects of anatomical confinement and boundary motions on the midgut dynamics, we numerically investigate the effect of removing these constraints from simulations. We specifically consider two orthogonal modifications: we consider the presence and absence of the confining walls defined in Sec. VI 7 by switching $W_1 = 0$ (top and bottom rows of Fig. S14), and consider relaxing the orientational boundary condition defined in Sec. VI 4 (left and right columns of Fig. S14). Throughout, we keep the end-points fixed to their experimental locations, as prescribed in Sec. VI 4.

Running these experiments with the optimal simulation parameters identified by the sweep, we find that with the viscoelastic update rules of the simulations protect the overall final shape to relaxing the orientation boundary conditions, while relaxing confinement leads to more dramatic morphological changes. Note that even in the absence of confinement, the resulting shape keeps a memory of the initial constrictions and is mostly unable to properly form the posterior domain.

#### VII. CANALIZATION OF MIDGUT CHIRALITY IN INTERPOLATED SHAPES

In Main Fig. 4, we explore how morphing initial conditions between representative wild-type and *byn>Myo1C* shapes lead to canalized chiral outcomes. In this section, we detail the construction of these interpolated shapes, and of self-contact, growth rates and confinement parameters. Other simulations parameters are as described in Table S 2.

##### 1. Initial condition interpolation and canalization

We use the SRV ("Square-Root Velocity") metric to interpolate between representative WT and *byn>Myo1C* initial conditions, allowing us to explore the effect of varying initial conditions in a restricted subspace of possible shapes [25]. The SRV representation $\mathbf{w}_\mathbf{x}(s)$ of a parametrized unit-length curve $\mathbf{x}(s)$ is given by

$$\mathbf{w}_\mathbf{x}(s) = \frac{\partial_s \mathbf{x}(s)}{\sqrt{|\partial_s \mathbf{x}(s)|}}, \tag{S56}$$

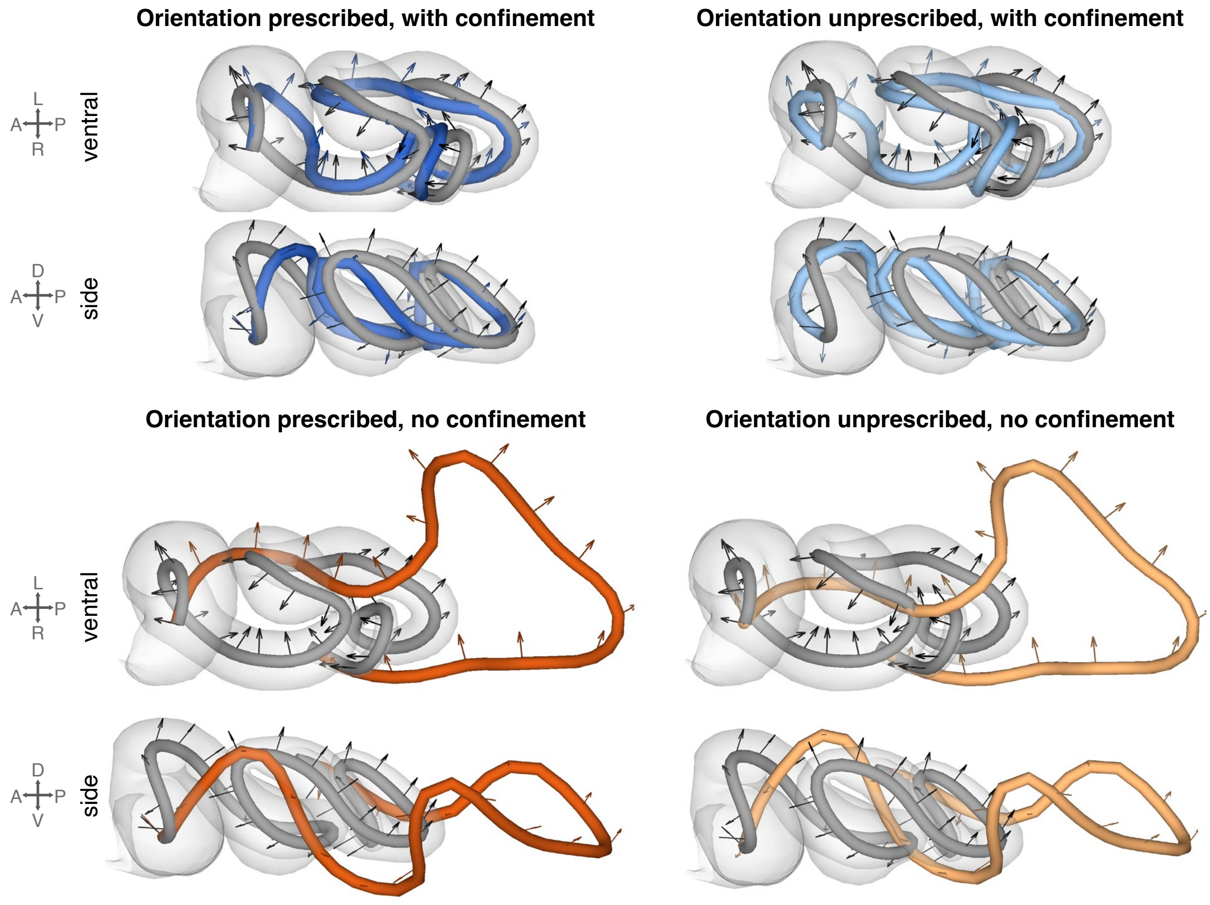


FIG. S14. Confinement and boundary orientation. Top row: simulations without prescribed frame boundary conditions. Bottom row: simulations without confining walls. Top left represents the conditions used throughout this study.

which can be inverted, starting from a known point $\mathbf{x}(0)$, as $\mathbf{x}(s) = \mathbf{x}(0) + \int_0^s \mathrm{d}p\, \mathbf{w}_\mathbf{x}(p)|\mathbf{w}_\mathbf{x}(p)|$. In this representation, we can define an inner product for a finitely sampled curve at $(s_i)_{i=1,\ldots,N+1}$

$$\langle \mathbf{w}_\mathbf{x}, \mathbf{w}_\mathbf{y} \rangle = \sum_{i=1}^{N} \mathbf{w}_\mathbf{x}(s_i) \cdot \mathbf{w}_\mathbf{y}(s_i) \tag{S57}$$

using the regular dot product in $\mathbb{R}^3$. Since $\langle \mathbf{w}_\mathbf{x}, \mathbf{w}_\mathbf{x} \rangle = 1$ for all unit-length curves $\mathbf{x}$, all unit-length curves live on the unit hypersphere $S^{3N-1} \subset \mathbb{R}^{3N}$. To interpolate between reference WT and *byn>Myo1C* centerlines $\mathbf{x}_{\mathrm{WT}}$ and $\mathbf{x}_{\mathrm{myo}}$, denoting $\mathbf{w}_{\mathrm{WT}}$ and $\mathbf{w}_{\mathrm{myo}}$ their SRV representations, we construct the curve $\mathbf{x}_\beta$, whose SRV representation is given by

$$\mathbf{w}_\beta = \frac{1}{\sin\theta} \left(\sin\left(\theta(1-\beta)\right) \mathbf{w}_{\mathrm{WT}} + \sin(\theta\beta) \mathbf{w}_{\mathrm{myo}}\right). \tag{S58}$$

Here $\theta$, with $\cos\theta = \langle \mathbf{w}_{\mathrm{WT}}, \mathbf{w}_{\mathrm{myo}} \rangle$, is the length of the geodesic on the hypersphere connecting the WT and *byn>Myo1C* curves in SRV space. By construction, $\mathbf{x}_{\beta=0} = \mathbf{x}_{\mathrm{WT}}$ and $\mathbf{x}_{\beta=1} = \mathbf{x}_{\mathrm{myo}}$; since this geodesic is a great circle on the hypersphere, we can also extrapolate in both directions beyond the reference samples by choosing $\beta \notin [0,1]$. We illustrate this procedure in Fig. S15a. In the main text, rather than $\beta$ we use $\alpha = 1 - 2\beta$, such that $\alpha = -1$ corresponds to WT while $\alpha = 1$ corresponds to *byn>Myo1C*.

Eq. (S58) allows us to interpolate the centerline position, but our simulations also require initial and boundary values for the material frame $Q$. To generate an interpolated frame $Q_\beta$, for all positions $s$ and at a given time $t$, we align the frame from $Q_{\mathrm{myo}}$ to the tangent $\mathbf{t}_{\mathrm{WT}}$, interpolate between WT and myo in the plane orthogonal to $\mathbf{t}_{\mathrm{WT}}$, before bringing back the resulting frame to be adapted to $\mathbf{t}_\beta$. Formally, defining $R_\mathbf{a}^\mathbf{b}$ as the minimal rotation sending vector $\mathbf{a}$ to $\mathbf{b}$, such that $\mathbf{b} = R_\mathbf{a}^\mathbf{b}\mathbf{a}$ and $\theta$ the angle about $\mathbf{t}_{\mathrm{WT}}$ between $Q_{\mathrm{WT}}$ and $R_1 Q_{\mathrm{myo}}$, the transform is defined

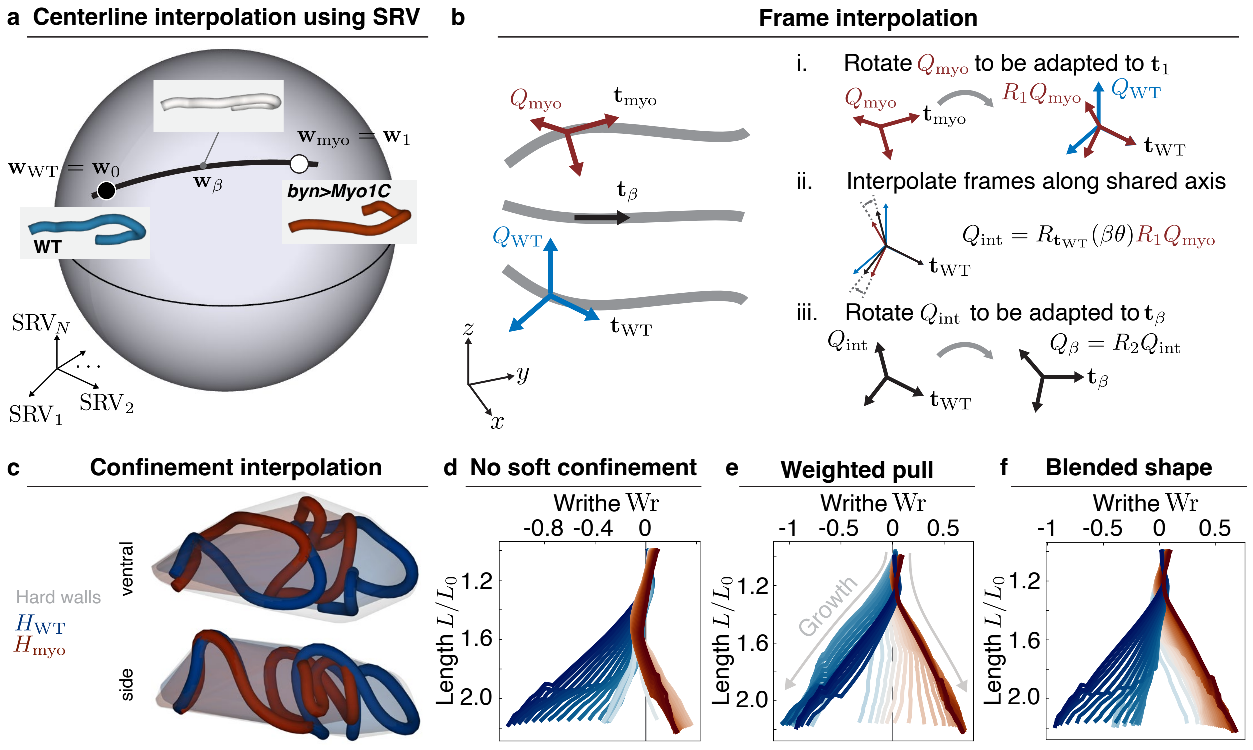


FIG. S15. SRV and frame interpolation. **a.** We use SRV interpolation to interpolate centerlines across conditions. **b.** Once centerlines are interpolated, we interpolate the frames in a shared adapted frame after minimally transporting them. **c.** Visualization of the hard confining walls (gray) and the two experimentally-derived convex hulls $H_{\rm WT}$ (blue) and $H_{\rm myo}$ (red). **d.** Wr vs. $L/L_0$ for interpolated simulations without soft confinement, that is only using the hard walls of panel **c**. **e.** Using the weighted pull scheme, introducing a soft confining force interpolating between the two experimental volumes. **f.** Using the 'blended shape' scheme, pulling within a single interpolated convex hull. All three schemes lead to canalized outcomes

through

$$Q_\beta = R_2 Q_{\rm int}, \quad \text{with } R_1 = R^{\mathbf{t}_{\rm WT}}_{\mathbf{t}_{\rm myo}}, \quad Q_{\rm int} = R_{\mathbf{t}_{\rm WT}}(\beta\theta) R_1 Q_{\rm myo}, \quad R_2 = R^{\mathbf{t}_\beta}_{\mathbf{t}_{\rm WT}} \tag{S59}$$

with $R_{\mathbf{t}}(\phi)$ the rotation of angle $\phi$ about vector $\mathbf{t}$. which is illustrated in Fig. S15b.

##### 2. Interpolation of growth rates

We also need to interpolate the growth rate $g(t,s)$, which we measure from the timeseries of interpolated centerlines by the change in length of each segment between consecutive frames (smoothed over frames), expressed in units of the initial length $L_0$. Each edge $e$ of the rod then elongates at $\dot{\ell}_e = w_e\, g(t, \bar{s}_e)$, where $s_e$ is its current fractional arclength and $w_e$ the fraction of an initial edge it carries after remeshing. Because the interpolation is performed in the square-root-velocity representation, intermediate curves are shorter than the two endpoint shapes, and the more so at late frames when the endpoint shapes differ most. As a result, the total growth of the input series dips in the middle of the sweep, so that without correction the final length is correlated with $\alpha$. To compare outcomes at equal length, we prescribe the total growth rate of every run while keeping its own pattern of growth along the rod and in time. Let $D(t)$ be the summed segment-length change of the input series per unit time (the total growth rate of the series). At every time step the edge rates are rescaled to

$$\dot{\ell}_e \;=\; w_e\, g(t, s_e)\, \frac{\Lambda(t)}{\sum_{e'} w_{e'}\, g(t, \bar{s}_e)}, \qquad \text{with } \Lambda(t) \;=\; (R-1)\, \frac{D(t)}{\int_0^T D(t')\,\mathrm{d}t'}, \tag{S60}$$

so that the total rest length grows as $L_0\big(1 + \int_0^t \Lambda\big)$ and reaches $R\,L_0$ at $t = T$ in every run; if the summed rate $\sum_{e'} w_{e'} g$ falls below $\Lambda/2$, a uniform rate per initial edge is added instead of rescaling. We take $R$ as the mean of the input growth of the two measured samples, $R = \frac{1}{2}\big[L(T)/L_0\big]_{\rm WT} + \frac{1}{2}\big[L(T)/L_0\big]_{\rm Myo}$. The final centerline lengths then agree to within 2% across the sweep, about 5% below $R\,L_0$ because the confined rod is slightly compressed relative to its rest length.

##### 3. Interpolation of self-contact and confinement

Interpolated simulations also require interpolating self-contact and confinement parameters. The former is straightforward: at each time point, the circumradii used for self-contact are linearly interpolated as $r_\beta(\bar{s}) = (1-\beta)r_{\rm WT}(\bar{s}) + \beta r_{\rm myo}(\bar{s})$. The latter is more subtle, and we consider three different schemes that reasonably interpolate between anatomical cavities of wild-type and inverted midguts.

First, we generate a 'hard wall', which contains all centerline points interpolated from the experimental samples, across all timepoints. This domain is generated from the convex hull of the union of all WT and *byn>Myo1C* data points for all $(\bar{s}, t)$, and across SRV-interpolated centerlines for all $\alpha$ (Fig. S15c). Alone, this already leads to canalized outcomes in the Writhe (Fig. S15d).

To examine the effect of interpolating the confinement between experimental settings, we also include an additional soft confinement that depends on the initial condition. Let $H_{\rm WT}$ and $H_{\rm myo}$ be the convex hulls of the WT ($\beta = 0$) and *byn>Myo1C* ($\beta = 1$) centerlines over all time points. A first 'Weighted pull' defines a average pulling force towards the two hulls: each node $i$ is subject to a force $\mathbf{f}_i$, with weights set by the interpolation parameter $\beta$:

$$\mathbf{f}_i = k_{\rm s}\Big[(1-\hat{\beta})\,\delta_i^{\rm WT}\,\hat{\mathbf{n}}_i^{\rm WT} + \hat{\beta}\,\delta_i^{\rm myo}\,\hat{\mathbf{n}}_i^{\rm myo}\Big], \qquad \hat{\beta} = \min(\max(\beta, 0), 1). \tag{S61}$$

Here, $\delta_i^H \geq 0$ is the distance of the node beyond the plane of the most violated face of hull $H$ ($\delta_i^H = 0$ inside $H$) and $\hat{\mathbf{n}}_i^H$ the inward normal of that face, as for the hard walls. The WT simulation ($\beta = 0$) is held in $H_{\rm WT}$, the *byn>Myo1C* simulation ($\beta = 1$) in $H_{\rm myo}$, intermediate simulations are pulled towards both, and extrapolated simulations ($\beta \notin [0,1]$) towards the hull of the nearer reference. We use $k_{\rm s} = 10^3$, ten times below the stiffness of the hard walls; with this value, the force with which the growing rod presses on the hard walls is balanced about $10\,\mu$m outside a hull, so that the rod can leave the hulls but is held near them. We find that the outcomes are still canalized (Fig. S15e), and this provides the confinement for the simulations shown in Main Text Fig. 4.

Finally, we also explore a 'blended shape' scheme where each simulation is softly confined to a single region that interpolates between the two hulls, defined by the weighted Minkowski sum of the two domains

$$C_\beta = (1-\hat{\beta})\,H_{\rm WT} \oplus \hat{\beta}\,H_{\rm myo} = \big\{(1-\hat{\beta})\,\mathbf{a} + \hat{\beta}\,\mathbf{b}\ :\ \mathbf{a} \in H_{\rm WT},\ \mathbf{b} \in H_{\rm myo}\big\}, \tag{S62}$$

a convex region that deforms continuously from $H_{\rm WT}$ at $\beta = 0$ to $H_{\rm myo}$ at $\beta = 1$ (and equals the hull of the nearer reference for $\beta \notin [0,1]$). $C_\beta$ is computed exactly as the convex hull of all pairwise sums $(1-\hat{\beta})\,\mathbf{v}_{\rm WT} + \hat{\beta}\,\mathbf{v}_{\rm myo}$ of the vertices of the two hulls. Every node outside $C_\beta$ is pushed back by $\mathbf{f}_i = k_{\rm s}\,\delta_i\,\hat{\mathbf{n}}_i$, with $\delta_i$ and $\hat{\mathbf{n}}_i$ the depth and inward normal of the most violated face of $C_\beta$, and the same $k_{\rm s} = 10^3$. The Writhe dynamics are qualitatively very similar to the ones from the simulations without interpolated soft confinement (Fig. S15f).

#### VIII. SCALING ARGUMENT FOR CONFINEMENT-DRIVEN CHIRALITY SELECTION

Simulations of the viscoelastic rod model of the midgut, started from initial shapes interpolated between WT and *byn>Myo1C*, show that the outcomes canalize into two mirror-image classes of final shapes, whose handedness follows the sign of the interpolation parameter $\alpha$. In this section, we develop a scaling argument arguing that this canalization is due to lateral confinement, which selects chiral over planar shapes. We discuss how the selected handedness depends on the initial shape and on remodeling, and compare to nonlinear simulations.

It is well-known that geometric effects are responsible for the sequence of sinusoidal-to-helical buckling of rods confined in tubes [26–28], such as drill strings and tubing in wells [29, 30]; a similar mechanism has also been proposed as a mechanism for heart tube looping [31]. We provide here a simple energetic argument to explain this sequence of instability.

We consider the midgut as an inextensible rod of length $L$ with bending stiffness $B_1$ and twisting stiffness $B_3$, whose ends are held at distance $L_{\rm e} < L$ along a fixed axis $\hat{\mathbf{z}}$, with fixed end tangents and frames. We obtain estimates of the bending and twisting costs of deformations, in the limit of small deflection and validate them against numerical simulations using the full model of Sec. VI in large deformations, finding agreement with the predicted phenomenology (Fig. S16).

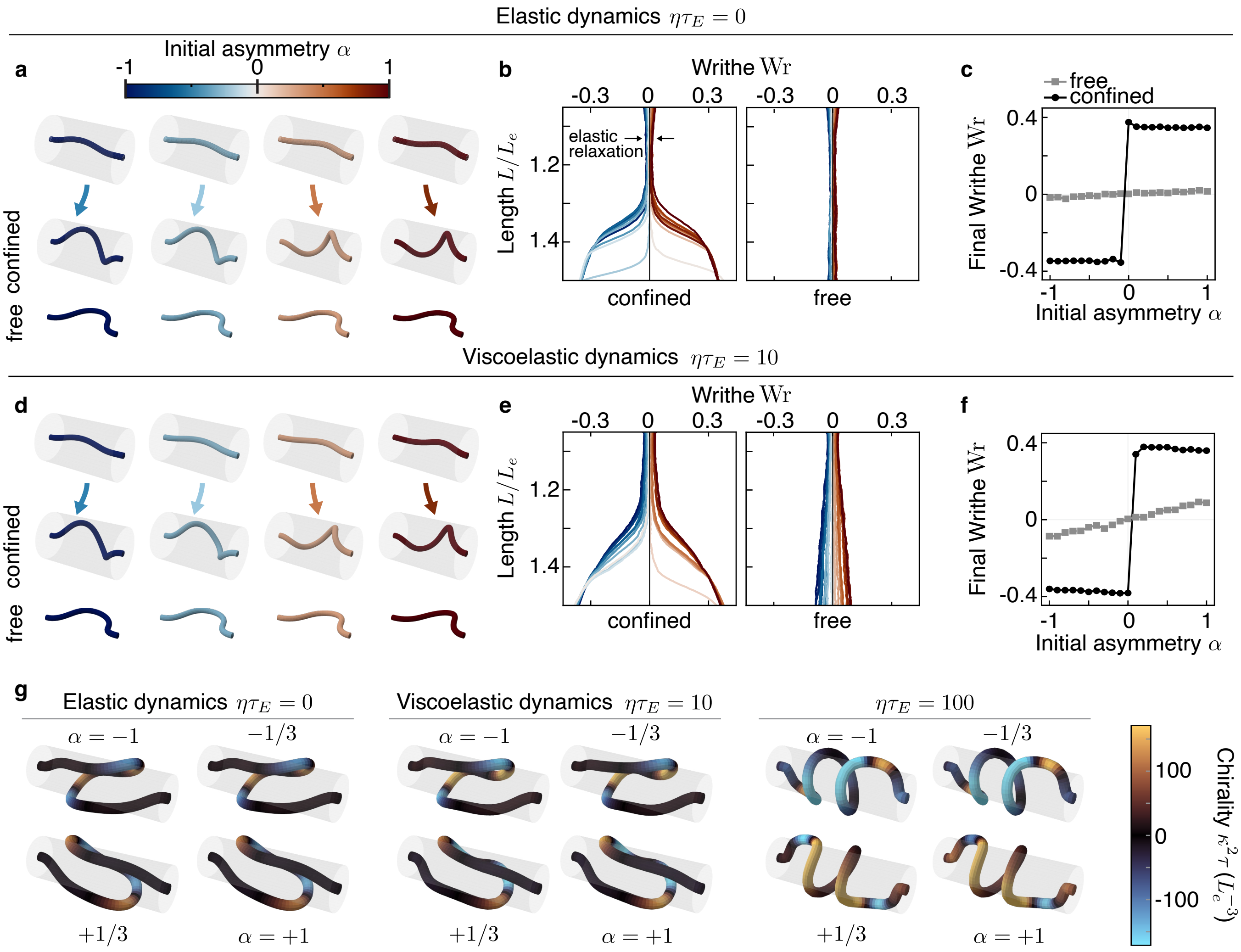


FIG. S16. Canalization of mirror-image helices by confinement. **a**. Initial shapes interpolated between left- and right-handed helices (colored by $\alpha$) grow from $L/L_e = 1.05$ to $1.3$ at rate $g\tau_E = 1$ either confined inside a channel of radius $R = 0.3L_e$ or unconfined (free) through elastic dynamics $\eta = 0$. $L_e$ is the distance between end points. **b.** With confinement, shapes canalize into two chiral shapes. In the absence of confinement, the initial handedness vanishes as the shape relaxes to planar. **c.** Quantification of the spread in Writhe as a function of the interpolation parameter $\alpha$. **d-f.** Same as above, but with viscoelastic dynamics $\eta\tau_E = 10$. The presence of viscoelasticity helps preserve the initial handedness even in the absence of confinement. All simulations use $B_3 = B_1$. **g.** At larger excess length, reducing twisting energy favors perverted configurations away from the fixed ends. This effect persists for weak remodeling ($\eta\tau_E = 10$), but vanishes at stronger remodeling ($\eta\tau_E = 100$), which then favors helices. The radius of the rod is arbitrary in visualizations.

For small deflections, we write the tangent as $\mathbf{t} \simeq (\psi_x, \psi_y, 1 - |\psi|^2/2)$ in a basis whose third vector is $\hat{\mathbf{z}}$, and collect its transverse components in the complex field $\psi = \psi_x + i\psi_y$. The transverse position of the centerline is $X(s) = \int_0^s \mathrm{d}s'\, \psi(s')$, the excess length is $\Delta = L - L_\mathrm{e} = \frac{1}{2}\int_0^L \mathrm{d}s\, |\psi|^2$, and the bending energy is $E_\mathrm{b} = \frac{B_1}{2}\int_0^L \mathrm{d}s\, |\psi'|^2$, with primes here denoting derivatives with respect to the arclength $s$.

In the absence of confinement, chiral shapes are disfavored. The bending energy splits into the energies of the two planar projections of the shape, $\frac{B_1}{2}\int \mathrm{d}s\, (\psi_x'^2 + \psi_y'^2)$, which does not depend on the relative phase of these projections and does not favor planar over chiral shapes. With fixed end frames, the link $\mathrm{Lk} = \mathrm{Tw} + \mathrm{Wr}$ is conserved as long as the rod does not pass through itself [17, 18]. For a rod that is initially straight and untwisted, $\mathrm{Lk} = 0$, so that any Writhe must be compensated by an equal and opposite Twist $\mathrm{Tw} = \frac{1}{2\pi}\int \mathrm{d}s\, \tau$. Thus, chiral shapes must carry Writhe which causes twist, which is penalized by twist rigidity. Consequently, in the absence of confinement, Euler buckling selects plane waves $\psi(s) = A\cos ks$.

Consider now a rod inside a hard tube of radius $\ell$, such that its transverse excursion is limited by the wall as $|X| \simeq \ell$, and compare two shapes of wavenumber $k$. On one hand, a planar wave with $\psi = A\cos ks$ reaches the wall for $A = k\ell$ and stores the excess length $\Delta = Lk^2\ell^2/4$. On the other hand, a helix with $\psi = Ae^{iks}$ uses both transverse directions: with the same amplitude $A = k\ell$, it stores twice as much arclength $\Delta = Lk^2\ell^2/2$. At fixed excess length, the helix can therefore buckle with a longer wavelength, $k^2 = 2\Delta/L\ell^2$ instead of $4\Delta/L\ell^2$. Since the bending energy of a single wave is $E_\mathrm{b} = B_1 k^2 \Delta$, this halves its bending cost,

$$E_\mathrm{b}^\mathrm{helix} = \frac{2B_1\Delta^2}{L\ell^2} = \frac{1}{2}\,E_\mathrm{b}^\mathrm{planar}. \tag{S63}$$

At small deflections, the Writhe is $\mathrm{Wr} = \frac{1}{4\pi}\int_0^L \mathrm{d}s\,\mathrm{Im}(\psi^*\psi')$ [17, 18], with $\mathrm{Wr} = 0$ for planar shapes. The helix thus has a uniform twist density $\tau = 2\pi\mathrm{Wr}/L = k^3\ell^2/2$, and the corresponding energy reads $E_\mathrm{tw} = \frac{B_3 L}{2}(2\pi\mathrm{Wr}/L)^2 = B_3\Delta^3/(L^2\ell^2)$. Compared to the reduction in bending energy, these geometric effects select helices when

$$\frac{E_\mathrm{tw}}{E_\mathrm{b}^\mathrm{planar} - E_\mathrm{b}^\mathrm{helix}} = \frac{B_3}{2B_1}\,\frac{\Delta}{L} < 1, \tag{S64}$$

which is the case for thin tubes where contact occurs at moderate excess lengths, unless the twist rigidity is very large. A wall that limits the transverse excursion therefore makes chiral shapes energetically favorable as soon as the rod grows to fill the available space.

When $\Delta \gtrsim L$, the argument above suggests that the rising twist costs make helices unstable. To reduce twist, the buckled shape can develop perversions [20, 31]: in this case, the twist cost no longer scales with $\Delta$, and chaining helices of alternating handedness still have lower bending energy than planar curves. More precisely, assume that helices and perversion have wavenumber $k$, and that a localized perversion has the same bending energy density as a helix. Then the bending cost from the perversion over a wavelength is $E_\mathrm{p} \propto B\kappa^2(1/k) = B\ell^2k^3$, where from above the curvature is $\kappa \sim \ell k^2$. Using the excess length $\Delta \propto Lk^2/\ell^2$, the ratio of twist energy to bending energy stored in the perversion is

$$E_\mathrm{p}/E_\mathrm{tw} \propto \left(\frac{B_1}{B_3}\right)\frac{1}{L\ell^2k^3} \propto \left(\frac{B_1}{B_3}\right)\left(\frac{\lambda}{\Delta}\right) \tag{S65}$$

with $\lambda \sim 1/k$ the wavelength. Thus, once $\Delta > (B_1/B_3)\lambda$, introducing a perversion is cheaper than twisting a single helix. Overall, geometric effects favor chiral shapes over planar ones, with perverted shapes favored at large deviation to reduce twisting energy.

A reflection of the shape maps $\mathrm{Wr} \to -\mathrm{Wr}$ and leaves both twisting and bending energies unchanged. In the absence of link, of a reference twist or of active torques, left- and right-handed shapes therefore have the same energy, and confinement selects one of two mirror-image states without preferring either. The final handedness is set by the sign of the Writhe when the rod reaches the confining boundary. The sign at that time is set by the initial configuration of the shape, provided elastic relaxation has not erased that initial asymmetry. The selection of chirality by confinement then explains the observed canalization into two mirror-image classes whose handedness follows the sign of $\alpha$ (Fig. S16a-f).

The selection relies on the elastic energy difference of Eq. (S63). Visco-elastic remodeling relaxes the reference curvatures towards the current ones at rate $\eta$, so that deformations are written into the rest shape and the corresponding elastic forces decay over a time $1/\eta$ [24]. If remodeling is too slow and the confining tube too large, the shape can relax to a planar state before hitting the confining walls, losing the memory of its initial state. If remodeling is too fast compared to growth, the elastic drive towards chiral shapes could vanish before it can act and noise could be integrated in the final shape [24]. Fast remodeling also cancels the twisting cost which favors perverted shapes over helices (Fig. S16g). Chiral canalization into perverted shapes therefore requires intermediate Fluidity $\eta\tau_E$, with $\tau_E$ the bending relaxation time, slow enough for twisting forces to persist over the growth period, and could explain why our parameter sweeps favor intermediate values of the fluidity (Sec. VI 8, Fig. S13).

#### IX. SUPPLEMENTARY VIDEO CAPTIONS

**Supplementary Video 1:** Overview of the pipeline used to reconstruct the three-dimensional geometry of the embryonic *Drosophila* midgut and extract its framed centerline. Multiview, multi-channel light-sheet microscopy captures the endoderm (blue) and circular visceral muscle (CVM) nuclei (yellow). The resulting volumetric images are segmented to obtain a three-dimensional surface mesh of the midgut, onto which the fluorescence data are mapped using `TubULAR`. The reconstructed surface is then parameterized by longitudinal ($u$) and azimuthal ($\phi$) coordinates. The positions of CVM nuclei, organized into four longitudinal curves along the organ, are used to construct an anatomically oriented material frame along the midgut centerline, as detailed in SI Sec. IV. This representation, consisting of the centerline $\mathbf{x}(s)$ and orientation frame $Q(s)$, provides the basis for quantifying midgut bending, twisting, and chirality.

**Supplementary Video 2:** Quantification of local organ chirality reveals *perverted* handedness. The unit vector tangent to the organ centerline defines a curve on the unit sphere as we sweep from anterior to posterior. The solid angle spanned by local portions of this curve is used to define a quasi-local measure of chirality, $q(s)$. Applying this measure to experimental samples, we find that the centerlines alternate handedness, thus exhibiting what is known as *perverted* chirality. See SI Sec. V for the definition and Fig. S10 for other experimental samples.

**Supplementary Video 3:** Simulations recapitulate coiling dynamics. Viscoelastic rod simulations are able to recover the post-constriction coiling of the midgut. Here, we show representative samples (WT 3, *byn>Myo1C* 1) from both ventral and side views, and show all 10 samples from ventral views. Colored tubes indicate simulations, and gray tubes indicate experimental centerlines.

**Supplementary Video 4:** Confinement canalizes chiral coils and induces perverted handedness. We simulate the growth of a family of viscoelastic rods with initial shapes interpolating between left-handed ($\alpha < 0$) to right-handed ($\alpha > 0$) chiral coils. In confined spaces, the interactions between wall and rod select for chiral shapes and focuses them to two distinct shapes. This focusing is absent without confinement. To mitigate twisting costs, the coil growing in confinement develops perversions, here shown for $\alpha = -1$ and measured using the quasi-local measure $q$ of Sec. V. In all simulations, $\eta\tau_E = 10$, and $g\tau_E = 1$, and the initial length is $L_0 = 1.05L_e$, with $L_e$ the distance between points. For confined runs, $R = 0.3L_e$.

**Supplementary Video 5:** Mechanical dynamics and interactions canalize organ shape. Morphing the wild-type initial shape into the *byn>Myo1C* initial shape, and beyond, along geodesics in an appropriate shape space [25] (SI Sec. VII, Fig. S15), the simulated centerlines cluster into two groups of opposite handedness (Fig. 4e,f, Supp. Video 5).


[1] Stuart Berg, Dominik Kutra, Thorben Kroeger, Christoph N. Straehle, Bernhard X. Kausler, Carsten Haubold, Martin Schiegg, Janez Ales, Thorsten Beier, Markus Rudy, Kemal Eren, Jaime I Cervantes, Buote Xu, Fynn Beuttenmueller, Adrian Wolny, Chong Zhang, Ullrich Koethe, Fred A. Hamprecht, and Anna Kreshuk. ilastik: interactive machine learning for (bio)image analysis. *Nature Methods*, 16(12):1226–1232, December 2019.

[2] Paolo Cignoni, Marco Callieri, Massimiliano Corsini, Matteo Dellepiane, Fabio Ganovelli, and Guido Ranzuglia. MeshLab: an Open-Source Mesh Processing Tool. In Vittorio Scarano, Rosario De Chiara, and Ugo Erra, editors, *Eurographics Italian Chapter Conference*. The Eurographics Association, 2008.

[11] Alain Goriely. *The Mathematics and Mechanics of Biological Growth*, volume 45 of *Interdisciplinary Applied Mathematics*. Springer New York, New York, NY, 2017.

[12] Basile Audoly and Yves Pomeau. *Elasticity and geometry: from hair curls to the non-linear response of shells.* Oxford University Press, Oxford, first published in paperback edition, 2018.

[24] Nicolas Romeo, David B Brückner, and Noah P Mitchell. Growth and remodeling control shape memory in morphogenetic rods. *arXiv preprint arXiv:2607.23907*, 2026.
